# The European Prevention of Alzheimer’s Dementia (EPAD) MRI Dataset and Processing Workflow

**DOI:** 10.1101/2021.09.29.462349

**Authors:** Luigi Lorenzini, Silvia Ingala, Alle Meije Wink, Joost PA Kuijer, Viktor Wottschel, Mathijs Dijsselhof, Carole H Sudre, Sven Haller, José Luis Molinuevo, Juan Domingo Gispert, David M Cash, David L Thomas, Sjoerd B Vos, Ferran Prados, Jan Petr, Robin Wolz, Alessandro Palombit, Adam J Schwarz, Chételat Gael, Pierre Payoux, Carol Di Perri, Joanna Wardlaw, Giovanni B Frisoni, Christopher Foley, Nick C Fox, Craig Ritchie, Cyril Pernet, Adam Waldman, Frederik Barkhof, Henk JMM Mutsaerts, for the EPAD consortium

## Abstract

The European Prevention of Alzheimer Dementia (EPAD) is a multi-center study that aims to characterize the preclinical and prodromal stages of Alzheimer’s Disease. The EPAD imaging dataset includes core (3D T1w, 3D FLAIR) and advanced (ASL, diffusion MRI, and resting-state fMRI) MRI sequences.

Here, we give an overview of the semi-automatic multimodal and multisite pipeline that we developed to curate, preprocess, quality control (QC), and compute image-derived phenotypes (IDPs) from the EPAD MRI dataset. This pipeline harmonizes DICOM data structure across sites and performs standardized MRI preprocessing steps. A semi-automated MRI QC procedure was implemented to visualize and flag MRI images next to site-specific distributions of QC features — i.e. metrics that represent image quality. The value of each of these QC features was evaluated through comparison with visual assessment and step-wise parameter selection based on logistic regression. IDPs were computed from 5 different MRI modalities and their sanity and potential clinical relevance were ascertained by assessing their relationship with biological markers of aging and dementia.

The EPAD v1500.0 data release encompassed core structural scans from 1356 participants 842 fMRI, 831 dMRI, and 858 ASL scans. From 1356 3D T1w images, we identified 17 images with poor quality and 61 with moderate quality. Five QC features — Signal to Noise Ratio (SNR), Contrast to Noise Ratio (CNR), Coefficient of Joint Variation (CJV), Foreground-Background energy Ratio (FBER), and Image Quality Rate (IQR) — were selected as the most informative on image quality by comparison with visual assessment. The multimodal IDPs showed greater impairment in associations with age and dementia biomarkers, demonstrating the potential of the dataset for future clinical analyses.

## 1. Introduction

In recent years, data sharing in neuroimaging research communities has become increasingly common, with multiple collaborative efforts for pooling data to form large, diverse samples (Thompson et al. 2014; Jack et al. 2008). Advantages of clinical multicenter imaging studies include obtaining larger samples of subjects from potentially diverse demographic populations, increasing statistical power, generalizability of sophisticated analyses (Friedman et al. 2006), and allowing the development of site- and scanner-independent imaging biomarkers (Pomponio et al., n.d.).

Research Data Management (RDM) is a complex and laborious process in large multisite neuroimaging studies (Nourani, Ayatollahi, and Dodaran 2019), requiring well-defined practice to ensure the accessibility and organization of data, and the provenance of processing steps (Borghi and Van Gulick 2018). The procedure of processing MRI data is highly flexible, and decisions made at early stages can lead to substantial variability in analysis outcomes (Carp 2012). Moreover, there is no standard systematic procedure for MRI quality control (QC) in large multicenter studies. While individual visual inspection is often too laborious, automated procedures can be promising, but highly dependent on study design and the type of sequences acquired (Esteban et al. 2017; Fidel Alfaro-Almagro et al. 2018).

The European Prevention of Alzheimer Dementia (EPAD) study is a prospective, multi-center, European cohort study that aims to characterize the prodromal stages of Alzheimer’s Disease (AD) and create a pool of well-characterized individuals for recruitment in potential pharmacological trials (Solomon et al. 2019). Multimodal imaging data are acquired at each center and centrally stored and processed.

This overview documents the methods and implementation details of the MRI data processing, QC procedures, and computation of several imaging-derived phenotypes (IDPs), as developed for the EPAD neuroimaging dataset. We then explored the sanity of IDPs by reporting their relationship with other biomarkers of neurodegeneration. The described pipeline is publicly available online (“https://github.com/ExploreASL/ExploreASL/tree/EPAD,” n.d.).

## 2. Methods

### 2.1 The European Prevention of Alzheimer’s Dementia Longitudinal Cohort Study (EPAD LCS)

EPAD eligibility criteria were age above 50 years and no history of dementia (clinical dementia rating (CDR) < 1). After providing written informed consent, participants underwent an extensive multimodal test battery including five outcome measurements: cognitive tests, demographics, cerebrospinal fluid (CSF) biomarkers, genetics, and brain MRI. Participants were followed up after 6 months, and after 12, 24, or 36 months, depending on their CDR score at baseline. Details on the EPAD rationale and study protocol are provided elsewhere (Solomon et al. 2019).

Here, we considered the EPAD LCS v1500.0 data release, which consists of the baseline data from the first 1500 participants included in the study.

### 2.2 The EPAD MRI acquisition protocol

The v1500.0 baseline data were acquired at 21 EPAD sites, including seven different scanner models from Siemens Healthineers, Philips Healthcare, and GE Healthcare. A common scanning protocol was developed during the preparation phase to keep between-site differences as small as possible while accommodating differences in scanner hardware and software limitations.

The EPAD LCS imaging protocol was composed of core and advanced sequences (Table 1). The core sequences provided structural information and confirmed participants’ eligibility status through baseline radiological assessment, and the advanced sequences were designed to investigate brain structure and function in greater detail. Whereas the core sequences were conducted at all sites (n=21), the advanced sequences were performed in a subset of EPAD sites (n=13) with 3T scanners.

**Table 1.**
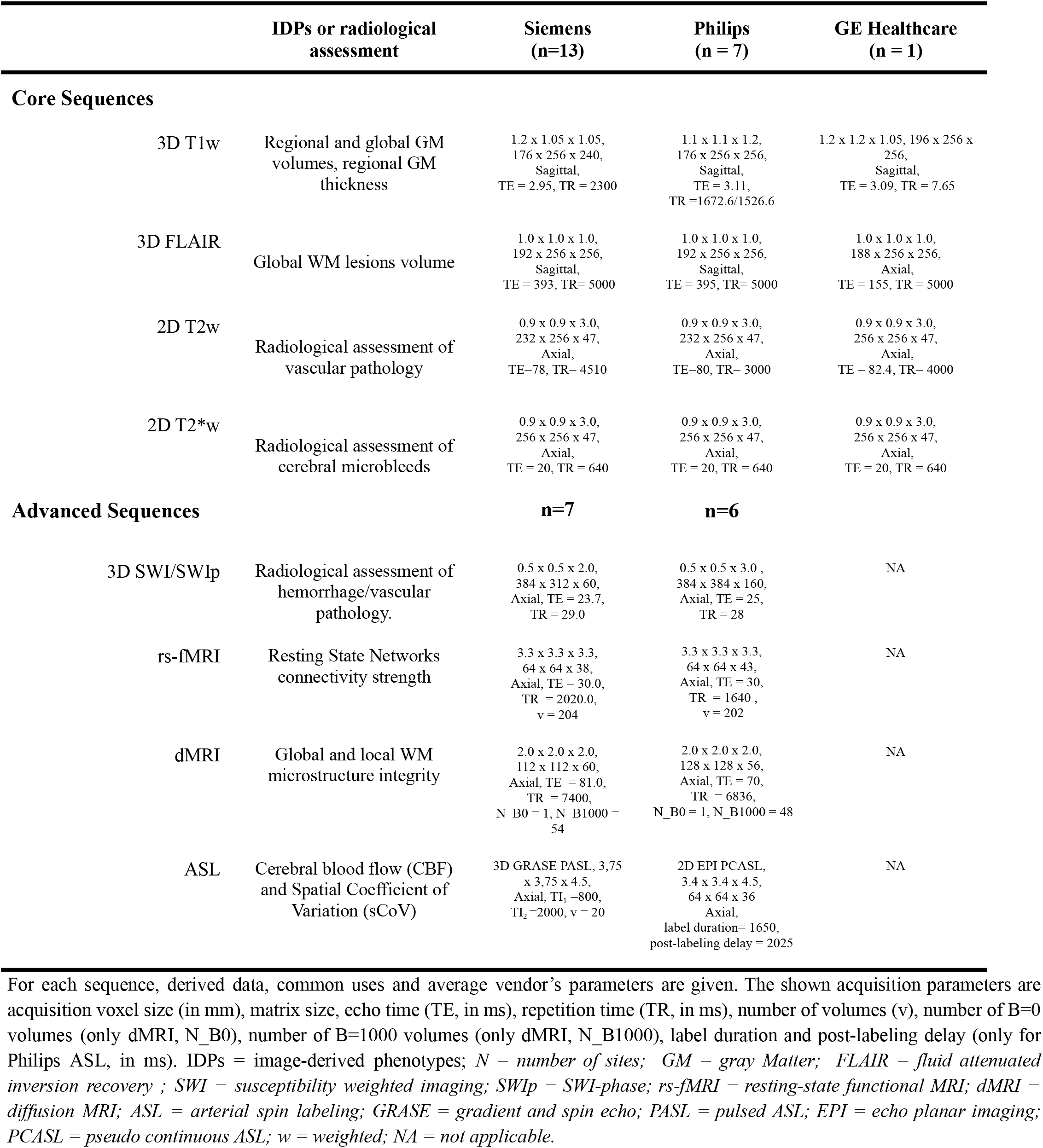
Core and Advanced MRI scan protocols.

### 2.3 The EPAD Imaging Pipeline

The EPAD image analysis pipeline consisted of four modules (Figure 1): 1) Curation of raw DICOM files, including harmonization of DICOM structure among sites, initial DICOM quality control (QC), and conversion to NIfTI; 2) Image preprocessing for core and advanced sequences; 3) Semi-automatic QC of processed data through an in-house code-based toolbox; 4) Computation of image-derived phenotypes (IDP) (Gong, Beckmann, and Smith 2020), i.e. extraction of numeric derivatives from images. Data sharing procedures are described in supplementary material section 2 and elsewhere (http://ep-ad.org/erap/).

**Figure 1.**
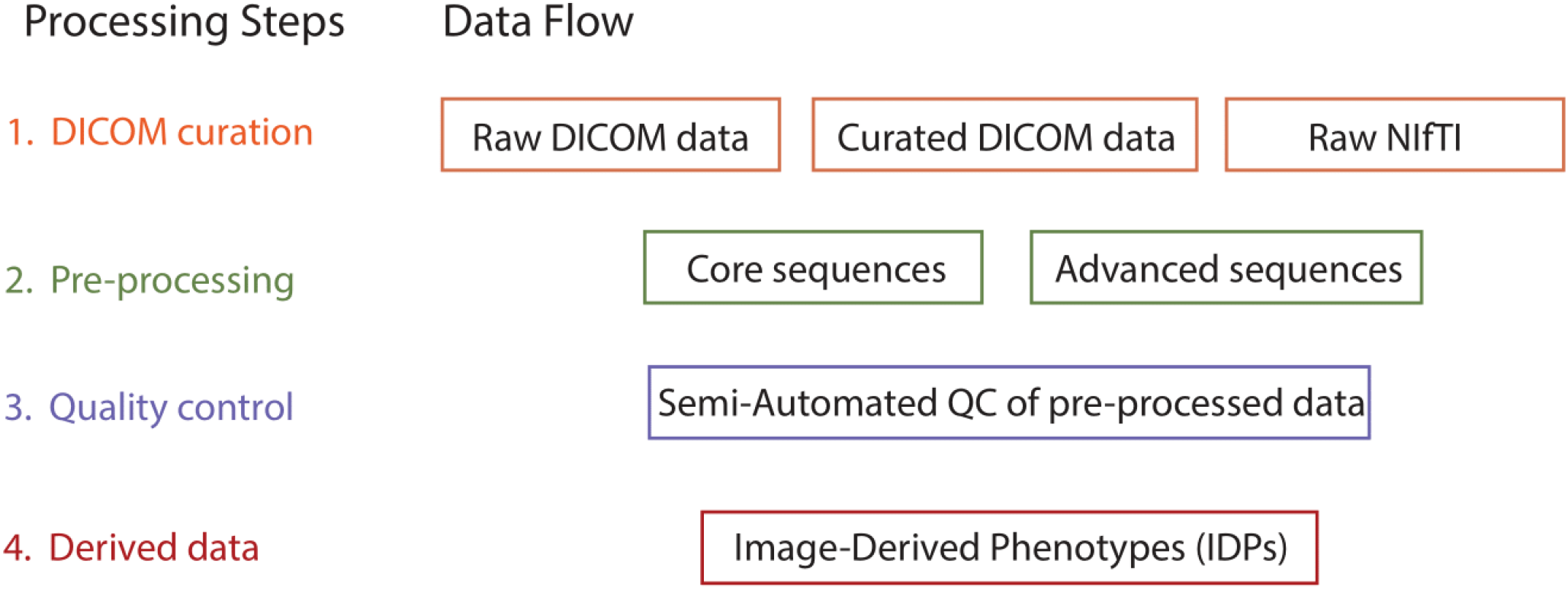
Image processing workflow in the EPAD study. *DICOM = Digital Imaging and Communications in Medicine; NIfTI = Neuroimaging Informatics Technology Initiative; QC = quality control; IDP = Image-derived phenotypes*.

All preprocessing steps were implemented in ExploreASL (Mutsaerts et al. 2020), an SPM-based toolbox designed to harmonize image processing for multi-center structural MRI and arterial spin labeling (ASL) studies. The toolbox was extended to include resting-state functional MRI (rs-fMRI) and diffusion MRI (dMRI) preprocessing routines based on SPM12 r7771 and FSL 6.0.2 (Penny et al. 2011; Jenkinson et al. 2012). Table S1 in supplementary material summarizes the main processing steps and their software implementations in the EPAD pipeline.

#### 2.3.1 DICOM Curation

EPAD data were initially collected in DICOM format in the */sourcedata/* folder and each subject was saved under its EPAD ID. DICOM headers were loaded with ExploreASL’s wrapper around DICOM ToolKit (DCMTK v 1.18) and DICOM fields SeriesDescription, or ProtocolName and ImageType were used to recognize the scan type using scanner-specific regular expressions provided in a TSV-file (Figure S1). The assorted zipped files and directories with DICOM files were sorted by scan type. DICOM header information was also used for DICOM QC, e.g., to verify that the StudyID is concordant across scans within an MRI session, to exclude duplicates, and verify completeness of DICOM series and consistency of its parameters.

Following the Brain Imaging Data Structure (BIDS) (Gorgolewski et al. 2016) convention, dcm2niiX r20190902 was used to convert DICOM images to NIfTI format (Li et al. 2016), along with an accompanying JavaScript Object Notation (JSON) sidecar storing relevant metadata. Additional modifications to convert the dcm2niiX output to BIDS then included: sorting and splitting SWI magnitude and phase NIfTIs, splitting the ADC image from the dMRI NIfTI, sorting the phase encoding polarity (PEPolar) scans, obtaining PEPolar parameters and adding them to the JSON sidecars, managing ASL-specific conversion issues (Clement et al. 2019), and manage vendor- and scanner-specific conversion issues.

#### 2.3.2 Image Preprocessing

An overview of the preprocessing steps for the core and advanced sequences is shown in Figure 2 and Table S1.

**Figure 2.**
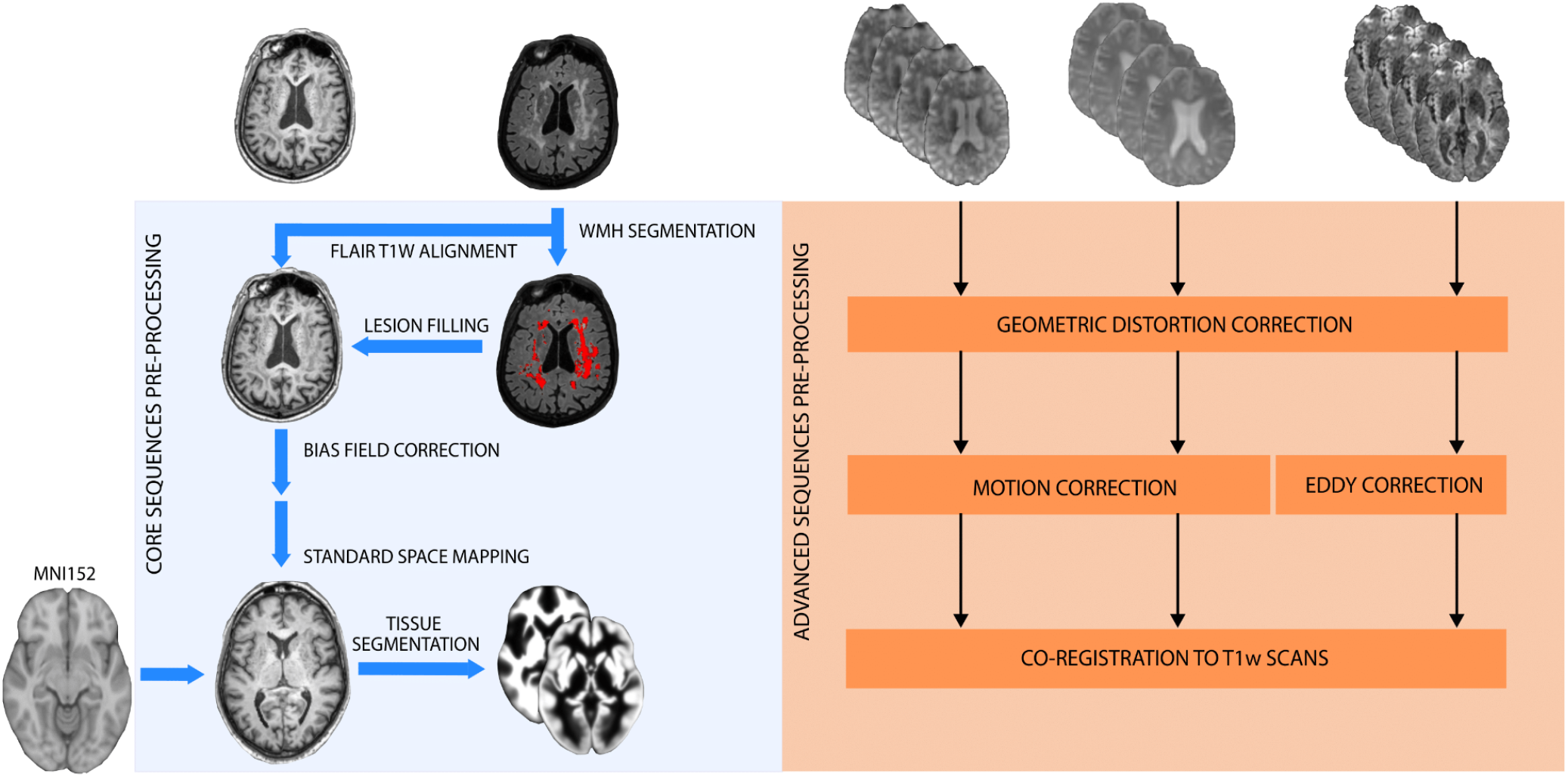
Schematic diagram of preprocessing steps. *Left:* the core sequences preprocessing pipeline performed on 3D T1w and 3D FLAIR scans; *Right:* the three common steps of the advanced sequences preprocessing pipeline. *Abbreviations: EPI = Echo-Planar Imaging*.

##### 2.3.2.1 Core Sequences Preprocessing Pipeline

The structural module of ExploreASL v1.0.2, described in (Mutsaerts et al. 2020), was used in combination with the Bayesian Model Selection (BaMoS) for WMH segmentation (Carole H. Sudre et al. 2015) to preprocess 3D T1 and 3D FLAIR images. Other 2D core sequences were only used for radiological assessment of patient eligibility and not preprocessed.

Preprocessing began with the registration of the 3D FLAIR to the 3D T1w (rigid-body) and the 3D T1w to the MNI center of mass (rigid-body). WMH segmentation was computed using BaMoS (Carole H. Sudre et al. 2015), a hierarchical unsupervised model selection framework simultaneously accounting for healthy tissue and unexpected observations. The resulting WMH segmentations were then used with the Lesion Segmentation Toolbox (LST) v2.0.15 (Gaser 2009) to fill these lesion areas on the T1w image, which can be present as hypointensities and affect subsequent segmentation and non-linear registration (Schmidt et al. 2019).

Tissue segmentation was then performed with the Computational Anatomy Toolbox (CAT) 12 (r1363), which estimates and corrects the bias field inhomogeneity in 3D T1w images, and iteratively improves the non-linear registration to MNI standard space and the creation of partial volume maps of gray matter (GM), white matter (WM) and CSF. All the described transformations were combined in a single transformation, to avoid multiple interpolations.

##### 2.3.2.2 Advanced Sequences Preprocessing Pipeline

Rs-fMRI, dMRI, and ASL were handled by different preprocessing submodules and underwent three common steps: geometric distortion correction, motion correction, and registration with the structural reference image T1w. Two additional steps were performed for dMRI (see below). SWI was only used for radiological assessment as no automated processing routines exist for this sequence.

###### A) Geometric Distortion Correction

Echo-planar images (EPI) suffered from geometric distortion due to B_0_ field inhomogeneity induced by magnetic susceptibility variability (Holland, Kuperman, and Dale 2010). For this reason, EPI scans were accompanied by short acquisitions with reversed phase-encoding gradient polarity, which have an opposite distortion pattern. From this pair of images, the geometric distortion was estimated using a previously described method (Andersson and Sotiropoulos 2016) as implemented in FSL topup (Stephen M. Smith et al. 2004), and used to correct the geometric distortion of the fMRI, dMRI, and ASL with a 2D EPI readout.

###### B) Motion Correction

Head motion within fMRI and ASL was estimated with rigid-body transformations using the SPM12 realign function (Friston et al. 1995), where the ASL images were combined with the threshold-free outlier exclusion method ENhancement of Automated BLood flow Estimates (ENABLE) (Mutsaerts et al. 2020; Shirzadi et al. 2018). In dMRI images, an additional off-resonance source is caused by the rapidly changing magnetic field inducing eddy currents (EC) within conductors (Zhuang et al. 2006). Head motion and eddy current-induced geometrical EPI distortions were estimated and corrected by the FSL Eddy tool (Andersson and Sotiropoulos 2016).

###### C) Structural registration

Advanced sequences were registered to the 3D T1w images using rigid-body transformations. Similar to the core preprocessing, all transformation fields are combined and applied simultaneously, to avoid multiple cumulative interpolations.

###### D) dMRI Tensor Fitting

In the dMRI preprocessing submodule the registration output was fed into the FSL Brain Extraction Toolbox (BET) (Stephen M. Smith 2002) and then into FSL DTIFIT, to fit the diffusion tensor model to the data and produce diffusion tensor imaging (DTI) scalars maps (fractional anisotropy (FA), and mean (MD), axial (AD) and radial (RD) diffusivity).

#### 2.3.3 Semi-automatic QC

To control the quality of the EPAD imaging cohort, we created an in-house workflow to perform semi-automatic QC of MRI data. This set of QC functionalities was written as an extension to ExploreASL called ExploreQC. The semi-automated QC procedure was based on two steps: feature estimation and visualization (Figure 3). ExploreQC code availability and software specifications are listed in section 4 of the supplementary material.

**Figure 3.**
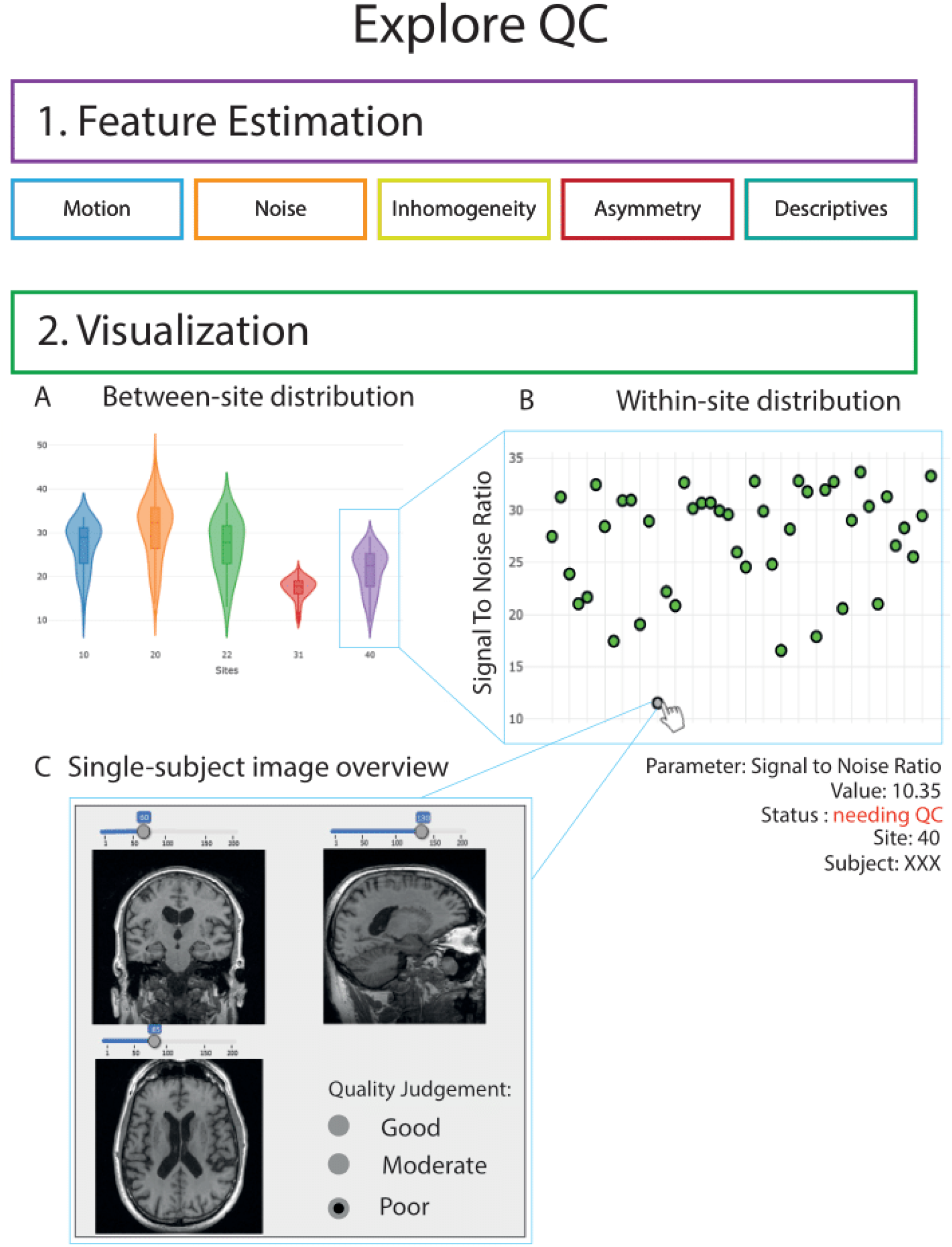
Overview of the quality control workflow. QC features are computed in the feature estimation module and cover 5 image features domains. Feature distributions can then be interactively inspected between-sites (2A) and within-sites (2B). Single-subject scans can be opened by clicking on the scatterplots (2C).

##### 2.3.3.1 QC - Feature Estimation

Image quality features were computed from five image feature domains: motion, noise, inhomogeneity, asymmetry, and descriptives, in line with recent MRI QC studies (Esteban et al. 2017; Fidel Alfaro-Almagro et al. 2018; Zarrar et al. 2015; Bastiani et al. 2019)). Definitions of domains and individual features are provided in table S2. All QC features were included in the MRI data release and can be referred to for study-specific inclusion/exclusion criteria.

##### 2.3.3.2 QC - Visualization

The visualization module consists of an interactive dashboard with violin and scatter plots for observing variation between and within sites, respectively (Figure 3). Individual scans can be visually inspected by selecting their data points on the scatter plots, allowing to visualize the scans themselves together with the QC features.

As a proof-of-concept for our semi-automated QC strategy, 3D T1w images were passed on to the visualization module. For all QC features, the difference with the site-specific mean was calculated as Z-score. Each scan was then sorted site-wise based on the sum of their absolute within-site Z-score values of all QC features. Scans with the 15% highest summed deviations were then automatically flagged for “needing visual QC”. To control for possible false-negative cases, an equivalent subset of non-flagged images was randomly selected and visually checked.

Visual QC was performed within the same pane by a single rater (LL), blinded to whether the image was an outlier or flagged as a random inlier. The image quality was categorized as “good” — desired MRI contrast visible and no artifacts or quality degradation detected, “moderate” — desired MRI contrast visible but some quality degradation, or “poor” — no desired MRI contrast visible and/or clear artifacts are present. These flags were added to the EPAD data release as a QC category advice for external researchers, together with the estimated features.

##### 2.3.3.3 QC Statistical Analysis

After visual inspection, we explored the association of the QC features with participants’ characteristics and with visual QC judgments. First, we used linear models to assess whether each QC feature distribution was related to the scanning site, age, sex, MMSE, amyloid, and APOE status of the participant. P-values were Bonferroni corrected.

To further investigate the features’ informative character in relation to the visual inspection, we built a logistic regression model with QC features as predictors of images showing quality issues among visual inspection. A stepwise backward parameter selection based on the Akaike Information Criteria was then performed to remove non-informative QC features from the final model.

#### 2.3.4 Image-derived phenotypes (IDPs)

IDPs are image-specific summary statistics that provide a quantitative way to investigate structural and functional brain characteristics (Gong, Beckmann, and Smith 2020).

##### 2.3.4.1 Core IDPs

Regional GM volume and cortical thickness are established phenotypes in neurodegenerative diseases (Scheltens et al. 2016). For 3D T1w sequences, we computed the volumetrics of several pipelines with different segmentation strategies. Template-based tissue volumetrics were computed from the CAT12-SPM tissue segmentation pipeline (Gaser 2009) described above. FreeSurfer v6.0.0 (Fischl 2012), one of the most widely used packages for measuring GM volumes and cortical thickness, was run on 3D T1w scans and included in the core IDPs release. A third method, the Learning Embeddings for Atlas Propagation (LEAP), was used to compute atlas-propagation-based regional volumes (Wolz et al. 2010). However, LEAP was run as a part of a separate and parallel pipeline and its results are not discussed here. WMH regional volumes were calculated from BaMoS segmentations and constitute 3D FLAIR derived data (C. H. Sudre et al. 2018).

##### 2.3.4.2 Advanced IDPs

###### rs-fMRI

Temporally correlated low-frequency (<0.1Hz) fluctuations in the rsfMRI signal are defined as functional resting-state networks (RSNs) (S. M. Smith et al. 2009). To identify RSNs in fMRI time series, a group-level independent component analysis (ICA) was performed by FSL Melodic (Beckmann and Smith 2004) on the preprocessed fMRI datasets. Two atlases of RSNs with a predefined number of independent components were generated: a low dimensional atlas with 20 and a higher dimensional atlas with 50 independent components. A dual regression approach was then used to obtain subject-specific RSNs (Nickerson et al. 2017). First, each RSN’s summary time course was estimated at the participant level by spatial regression of the full set of independent components from the high and low dimensional Melodic analysis against each participant’s fMRI data. Second, the resulting time courses were regressed into the same participants’ fMRI data to obtain subject-specific RSN maps. fMRI IDPs were computed as the mean within-network connectivity strength per subject.

###### dMRI

Tract-based spatial statistics (TBSS) is an automated, observer-independent approach for assessing voxel-wise fractional anisotropy in white matter tracts across groups of dMRI scans (Stephen M. Smith et al. 2006). The brain-extracted fractional anisotropy (FA) images, obtained after tensor fitting, were aligned into a common space using nonlinear registration. Next, the mean FA image was thinned to create a mean FA skeleton representing the center of all tracts common to the group and use it as a mask to compute individual FA values. Diffusion MRI IDPs were computed as global and regional FA features from the JHU ICBM-DTI-81 atlas (Wakana et al. 2004).

###### ASL

Arterial spin labeling perfusion MRI acquires cerebral perfusion *in vivo* in a non-invasive manner. Recent findings have shown that the 2D EPI readout on previous Philips software releases exhibits fat-saturation-related artifacts that considerably alter the quality of these scans, even to the point of being unusable (Mutsaerts et al. 2020). Therefore, we derived data only on a subset of ASL images that did not suffer from this artifact. Mean cerebral blood flow (CBF) and spatial coefficient-of-variation were computed as described in (Mutsaerts et al. 2020).

##### 2.3.4.3 IDPs relationship with AD markers

We finally explored the sanity and relevance of IDPs by assessing their relationship with age and other non-imaging data whose association with brain phenotypes has been established in the Alzheimer literature. Gray matter volume association with age was assessed through the Pearson correlation coefficient of both global and regional values. Increase in volume of WMH per additional year of age was evaluated as previously described (C. H. Sudre et al. 2018) in males and females separately, to account for sex differences in cerebrovascular disease (Haberman, Capildeo, and Rose 1981). As nonlinear trajectories of functional connectivity have been observed with age (Persson et al. 2014), mean within RSN connectivity quadratic relationship with age and differences between CDR = 0 and CDR = 0.5 participants were evaluated. dMRI and ASL IDPs correlation with age was computed, interacting age with amyloid status as defined in (Ingala et al. 2021) and Apolipoprotein E (APOE) e4 allele carrier status, respectively. As an initial exploratory data analysis, we here only report several preliminary associations. We did not explore the statistical significance level or effect of covariate correction on these correlations.

## 3. Results

### 3.1 The EPAD LCS Baseline Imaging Dataset

Of the 1500 screened participants, 144 did not fulfill the EPAD LCS eligibility criteria and were excluded from the analysis (Solomon et al. 2019). The resulting dataset has a complete set of core sequences for 1356 participants. Advanced sequences were performed at thirteen sites, with 756 SWI, 842 fMRI, 831 dMRI, and 858 ASL scans acquired. The mean age was 65.46 ± 7.14 and 775 (55.71 %) were female. All participants were without dementia at inclusion with an average MMSE of 28.58 ± 1.66, and 259 (19.11%) had a CDR of 0.5. A total of 1246 participants had CSF measurements, of which 845 (67.8 %) individuals were CSF amyloid negative and 401 (32.2 %) CSF amyloid positive, following previously defined study-specific cutoffs (Ingala et al. 2021). A detailed description of the baseline clinical and demographic characteristics of the EPAD cohort can be found in (Ingala et al. 2021).

Pre-processing steps were successful for all the T1w images, while failed for 8.6 % of fMRI and 8.6% of dMRI data, mostly due to high head motion or lack of necessary files (e.g. reverse-phase). A high percentage (62.3%) of the ASL sequences showed fat-saturation artifacts and are currently excluded from the dataset; preprocessing failure rate for the remaining ASL sequences was 0%. The final preprocessed EPAD dataset resulted in 1356 T1w and FLAIR, 770 fMRI, 759 dMRI, and 237 ASL scans. Details on the number of scanned and available processed data are provided in Figure 4.

**Figure 4.**
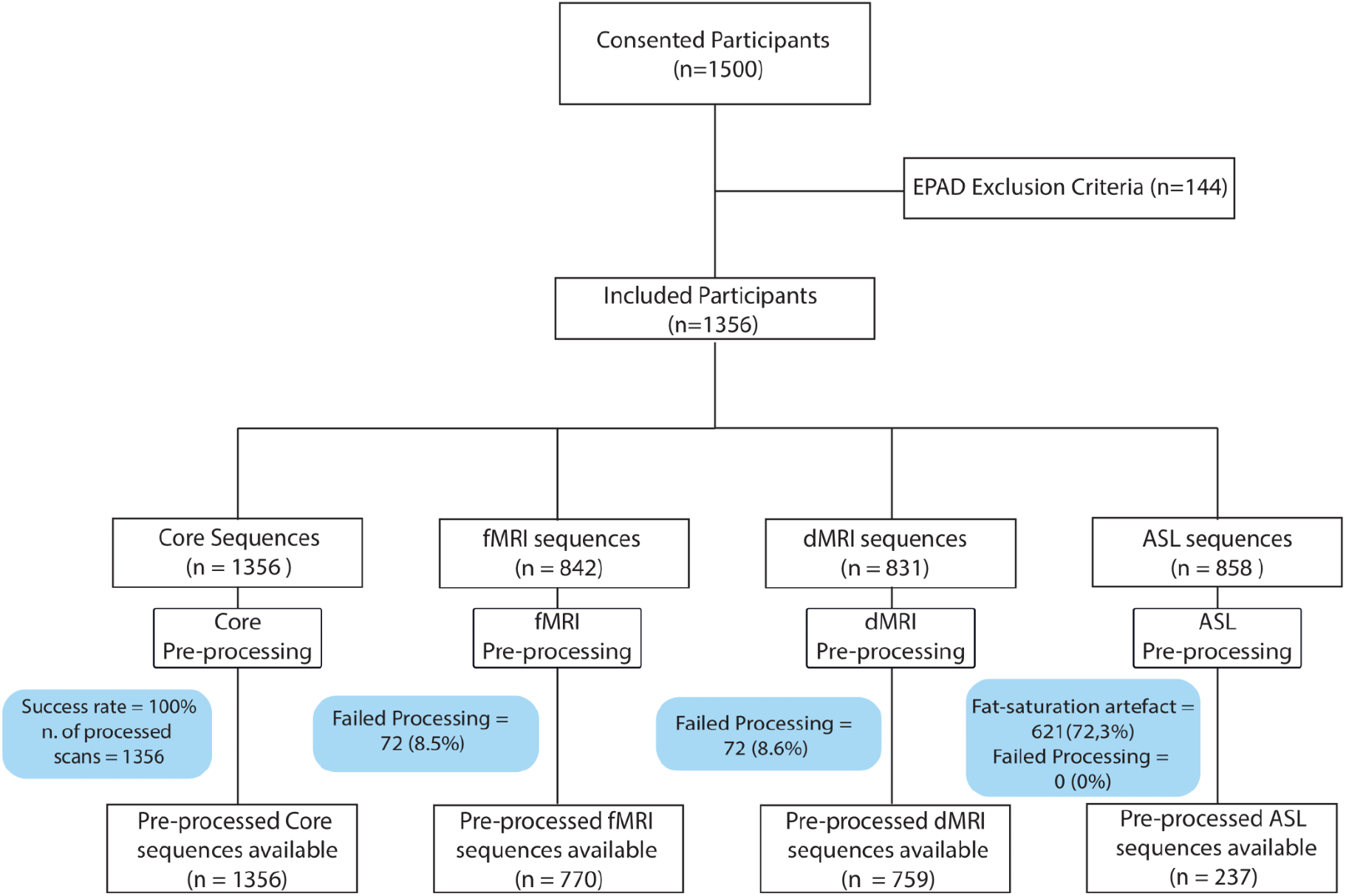
Consort diagram representing number of scanned and successfully processed sequences. Abbreviations: EPAD = European Prevention of Alzheimer’s Dementia; T1w = T1 weighted; FLAIR = Fluid attenuated inversion recovery; fMRI = functional magnetic resonance imaging; dMRI = diffusion magnetic resonance imaging; ASL = Arterial spin labeling.

### 3.2 Quality Control

49 QC features from 3 different image modalities (3D T1w = 12, fMRI = 17, dMRI = 20) were computed and are included in the current release.

Linear models showed that the QC feature distribution of T1w, fMRI, and DTI significantly differed between scanning sites. Participants’ demographic characteristics, such as age and sex, were related to the motion- and noise-related QC features for the 3D T1w and fMRI images (P<0.05), but not for the dMRI images (P>0.05). Amyloid status and MMSE were associated with 3D T1w motion- and noise-related QC features. APOE ε4 status was not related to any QC feature (Table S3).

Based on within-site distributions of the 12 QC features extracted from 3D T1w images, 197 scans (15% of the whole sample) were flagged as “needing-QC’’ for the visualization module. Of those, 16 were categorized as “poor quality” on visual inspection, while 51 were labeled as “moderate quality”. In the same number of visually inspected non-flagged scans, only one was labeled “poor” while 10 were judged to have “moderate” quality.

In the stepwise logistic regression analysis, backward parameter elimination removed four out of the 12 structural QC features in the final reduced model (Table 3). Five of those — Signal to Noise Ratio (SNR), Contrast to Noise Ratio (CNR), Coefficient of Joint Variation (CJV), Foreground-Background energy Ratio (FBER), and Image Quality Rate (IQR) — were significantly associated with the QC visual judgment and thus considered the most informative. When comparing the reduced model (i.e., after backward elimination) with the initial full model (i.e. before backward elimination), there was no significant loss of fitting. Adding site, sex, and age as covariates in the model did not significantly affect our findings (data not shown). A detailed description of the presented QC parameters can be found in section 5 of the supplementary material.

**Table 3.** Results of stepwise backward parameter elimination in QC logistic regression.

| *Parameter* | *Domain* | *Estimates* | *CI* | *p* |
| --- | --- | --- | --- | --- |
| (Intercept) |  | 1.47 | 0.6014 – 2.3455 | **0.001** |
| Signal to Noise Ratio (SNR) | Noise | -0.015 | -0.0210 – -0.0091 | **<0.001** |
| Contrast to Noise Ratio (CNR) | Noise | -0.089 | -0.1246 – -0.0544 | **<0.001** |
| Coefficient of Joint Variation (CJV) | Inhomogeneity | -0.062 | -0.1160 – -0.0073 | **0.026** |
| Foreground-Background energy Ratio (FBER) | Inhomogeneity/Motion | 0.003 | 0.0013 – 0.0045 | **<0.001** |
| Asymmetry Index percentage (AI_perc) | Asymmetry | -0.001 | -0.0013 – 0.0001 | 0.107 |
| Image Quality Rate (IQR) | Inhomogeneity/Noise | 0.317 | 0.2660 – 0.3675 | **<0.001** |
| Kurtosis in the CSF (CSF_k) | Descriptives | -0.014 | -0.0295 – 0.0022 | 0.092 |
| White Matter to Maximum Intensity ratio (WM2MAX) | Inhomogeneity/Descriptives | -0.761 | -1.7109 – 0.1890 | 0.116 |

The reduced model included 8 parameters, 5 of which showed a p-value of less than 0.05 in the association with the visual QC judgment. Estimates represent the change in the log odds of the outcome for a one-unit increase in the predictor variable. P-values < 0.05 are shown in bold

### 3.3 Image-derived phenotypes

358 IDPs per subject were computed from core sequences and provided information about total and regional GM volumes, cortical thickness, and white matter lesions. Example 3D T1w-IDPs are shown to correlate with age in Figure 5 and Figure S2. The FreeSurfer results showed negative associations of regional GM volumes with age, most markedly in the hippocampus (r=-0.34). A similar association is shown for the global GM volume computed with the CAT12 segmentation. From the set of 36 regional WMH volumes quantified on the 3D FLAIR scans, we found a frontal and parietal prevalence of WMH increase with aging, with no clear differences between sexes (Figure 6).

**Figure 5.**
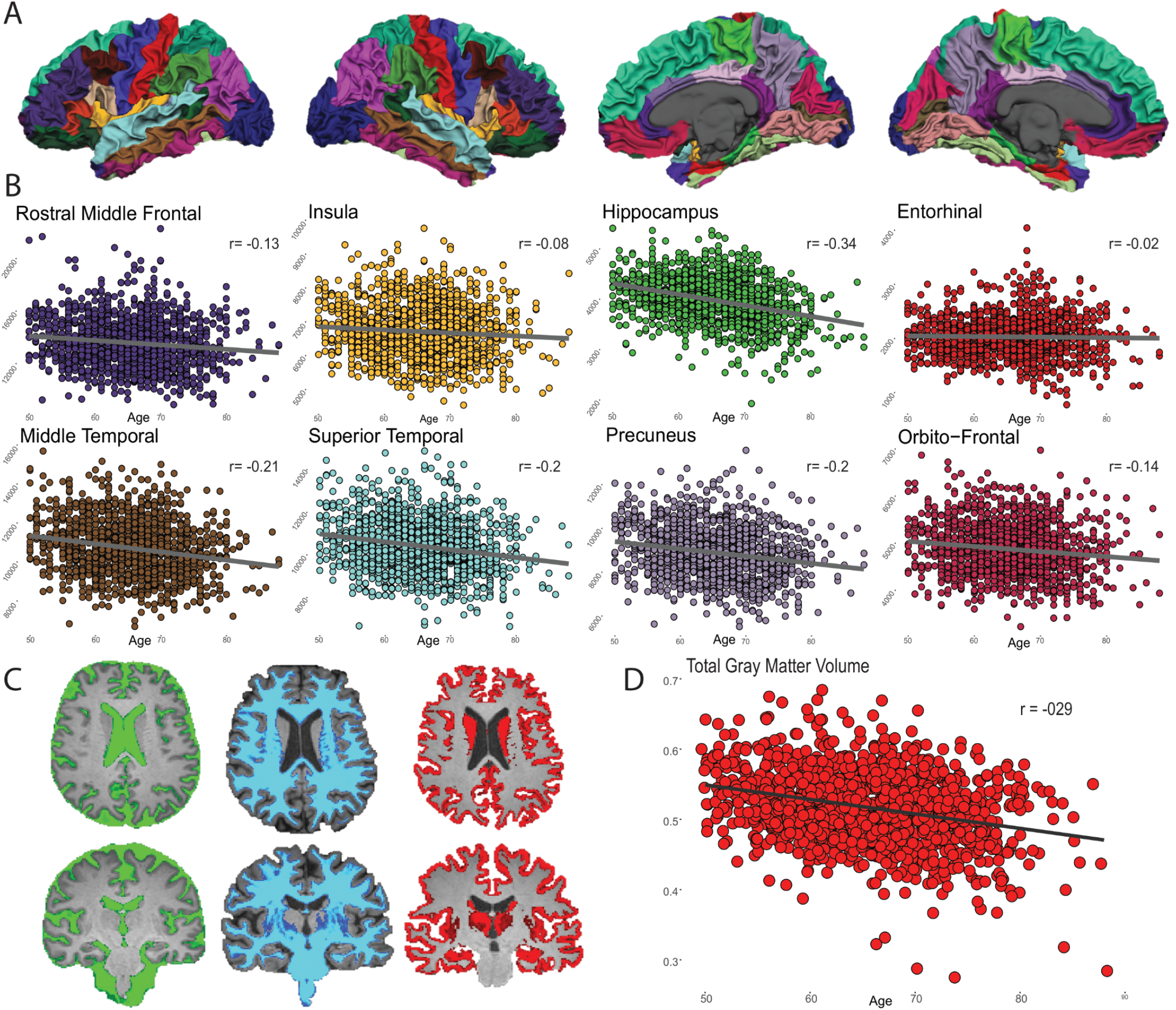
3D T1w derived phenotypes. Example association between core sequences derived data and age. A) FreeSurfer surface reconstruction of one 3D T1w image; B) Association of eight cortical regional volumes with age; C) CAT12 tissue segmentation of one 3D T1 scan output: green = cerebrospinal fluid, blue = white matter, red = gray matter; D) Association of total gray matter volume, as computed with CAT12 segmentation, with age.

**Figure 6.**
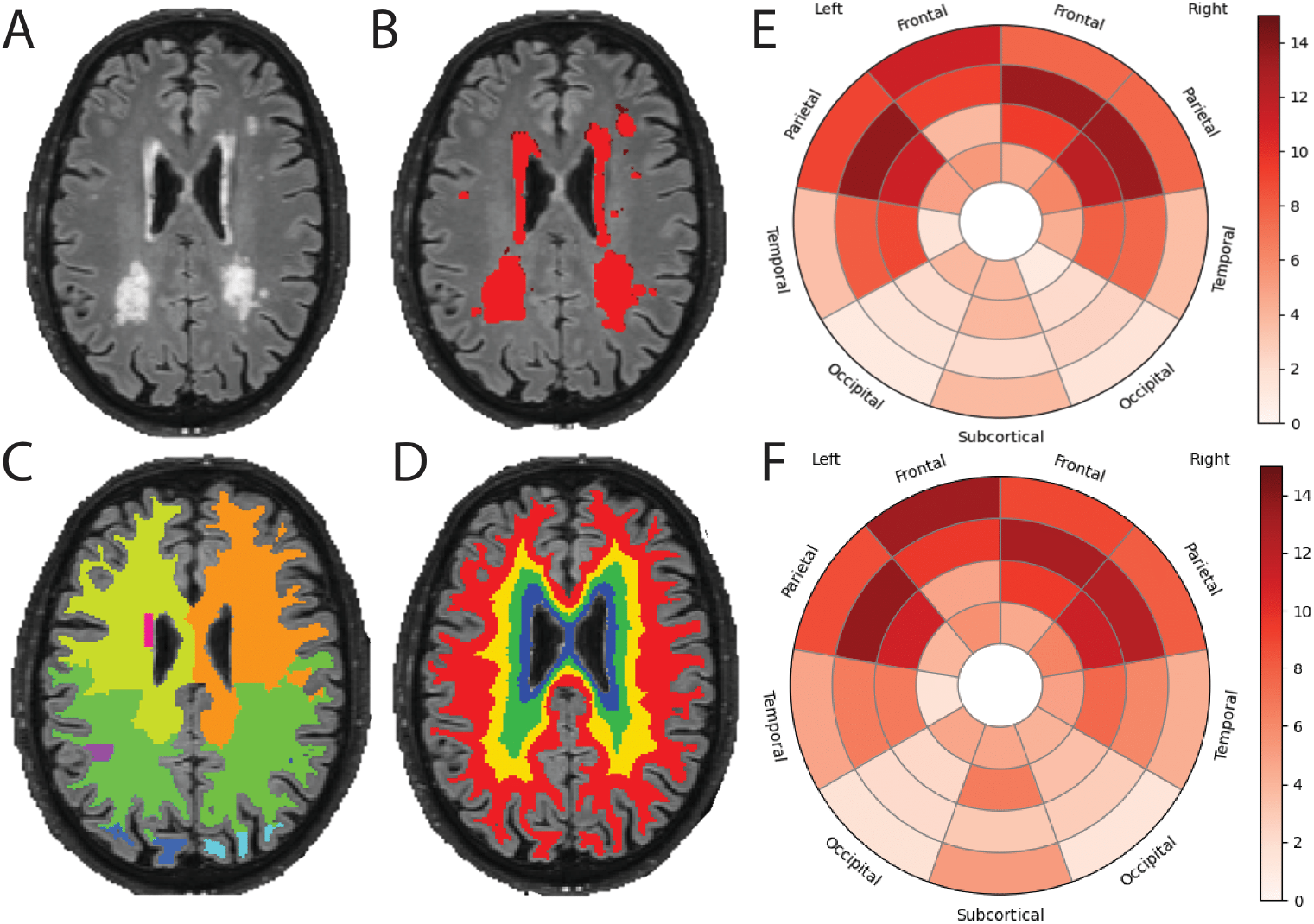
FLAIR derived phenotypes. A) Example FLAIR scan from one EPAD participant with relatively high lesion volume; B) Result of the white matter hyperintensities (WMH) segmentation using BaMoS; C, D) Lobes and layer atlases, respectively, used for regional WMH volume computation, methodological details are given in (C. H. Sudre et al. 2018); E, F) Effect of age on WMH frequency (expressed in percentage of increase in frequency, i.e. the proportion of lesion in a given region, per additional year of age) for male and female respectively.

The mean network connectivity was computed for 70 ICA components for the low and high dimensional ICA resulting in 70 fMRI IDPs. Figure 7A shows six RSNs commonly found in the literature (S. M. Smith et al. 2009), identified in the low-dimensional ICA. Complete sets of extracted components from both high and low dimensional ICA are shown in supplementary material Figure S3 and S4. RSN’s non-linear relationship with age and cognitive status was assessed and is shown in Figure 7B. Non-linear relationship of mean within-network connectivity with age differed by CDR groups, more strongly in the default mode and visual networks.

**Figure 7.**
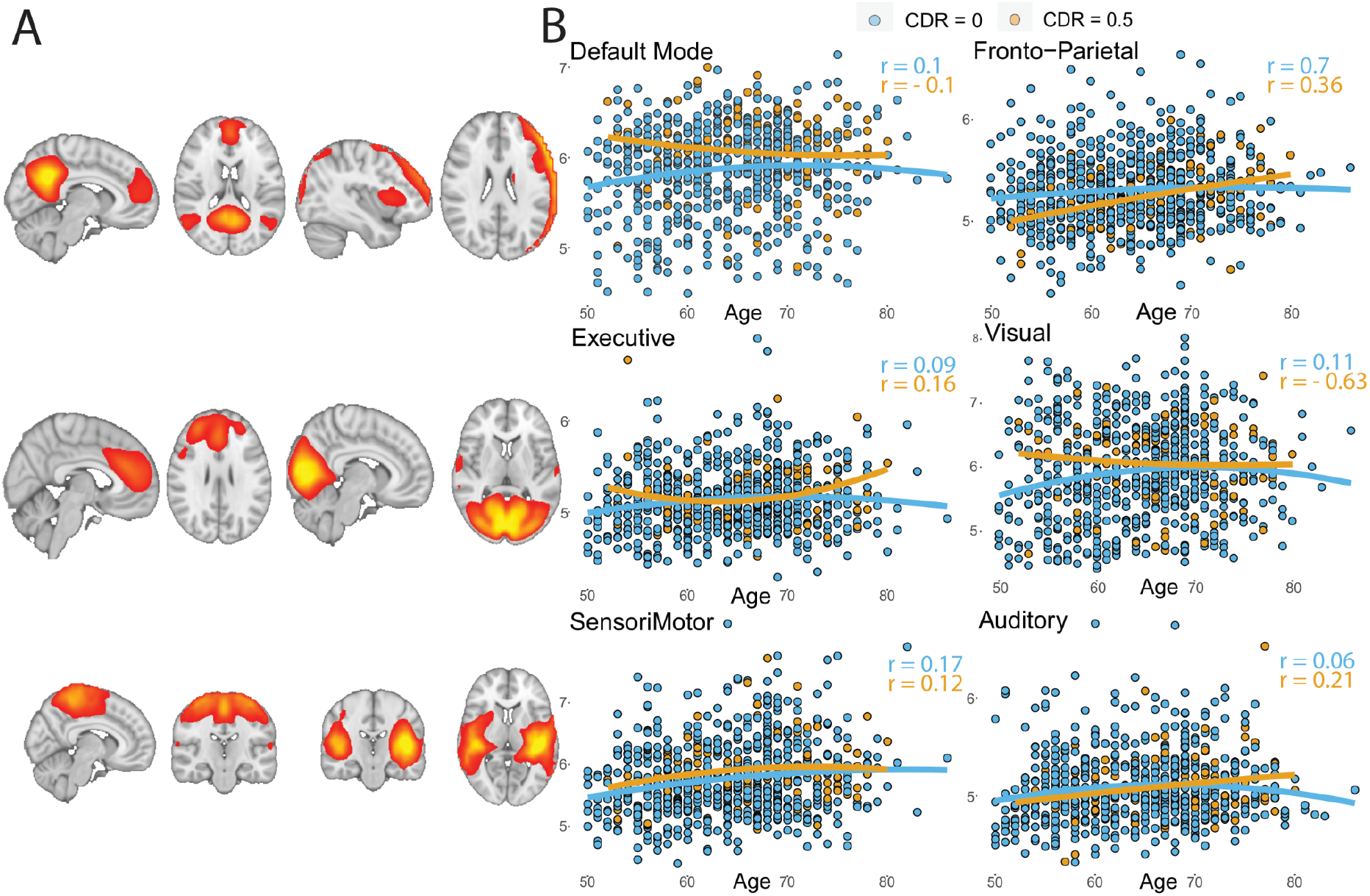
Resting-state fMRI derived phenotypes. A) Six group resting-state networks spatial maps from a low dimensional (20 independent components) melodic ICA; B) Scatter plots showing the non-linear relationship of mean within-network functional connectivity with age, grouped by clinical dementia rating (CDR) score. R values are computed as the Pearson correlation coefficients between the quadratic age term (age^2^) and mean within network connectivity values.

Global and local skeletonized FA values for the 48 regions from the JHU ICBM-DTI-81 atlas (Wakana et al. 2004) were calculated as dMRI IDP. Negative correlations of age with both global and regional WM integrity were found, with more negative coefficients in the amyloid positive participants (Figure 8).

**Figure 8.**
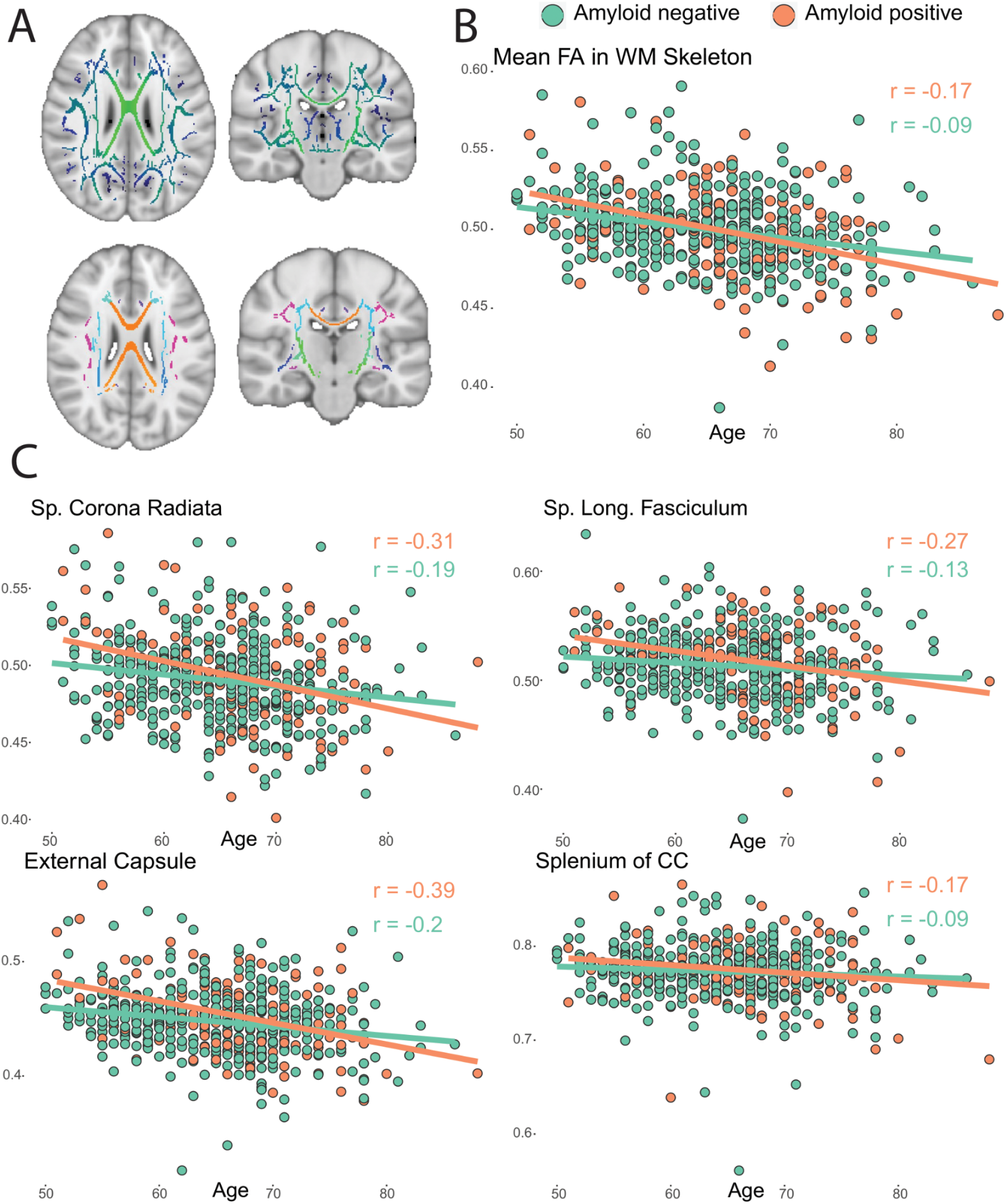
Diffusion MRI-derived phenotypes. A) The FA skeleton as computed in the TBSS pipeline (upper row) and the skeletonized white matter atlas used to extract local FA values (bottom row); B) Association of mean global FA values with age and amyloid status (as defined in (Ingala et al. 2021)). C) Association of 4 regional FA values with age and amyloid status. *Abbreviations: FA = Fractional anisotropy; TBSS = Tract based spatial statistics; WM = White Matter; Sp. = Superior; CC = Corpus Callosum; r = Pearson correlation coefficient*.

Usable ASL data (without fat-saturation artifact) were derived from 237 participants. Mean CBF maps across participants showed considerable regional variability across the brain with high perfusion in the cingulate and precuneus GM and lower perfusion in basal ganglia (Figure 9A). An atypical relationship between CBF and age was observed in APOE-e4 allele carriers, showing a lack of CBF decline with age. In contrast, non-carrier participants showed a reduction of CBF with advancing age (Figure 9B).

**Figure 9.**
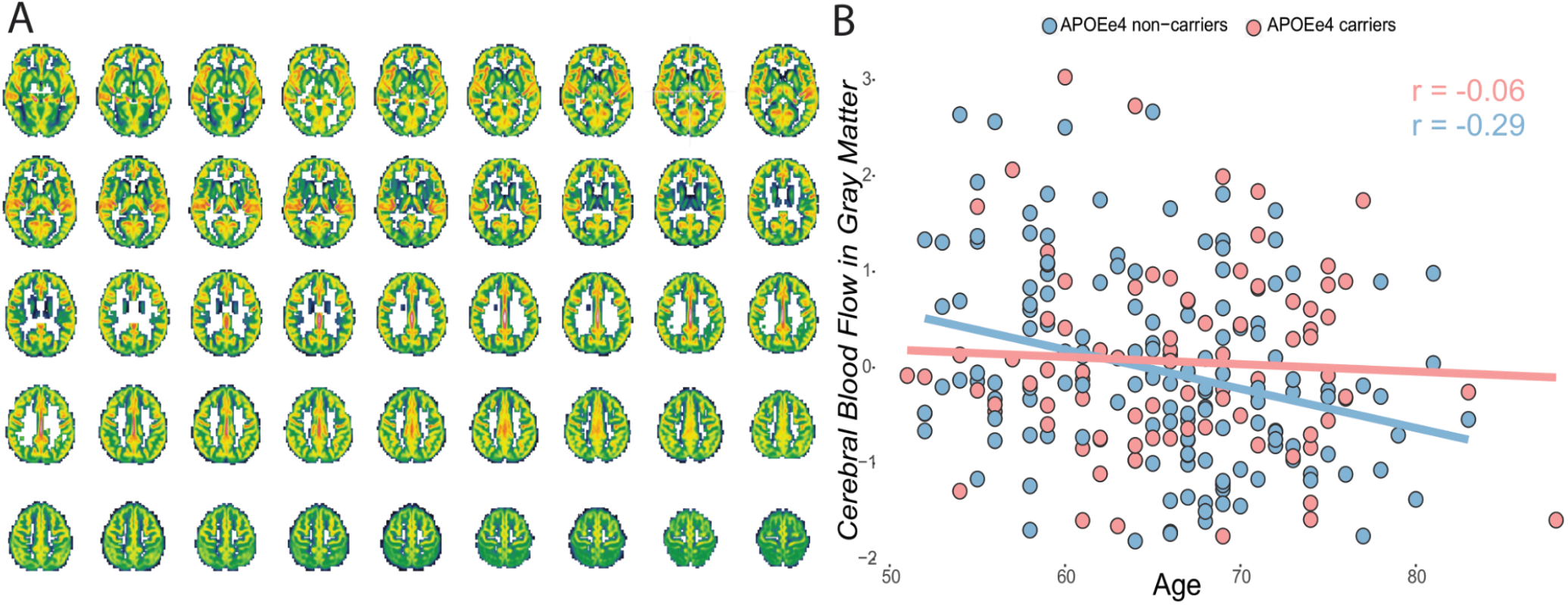
Arterial spin labeling IDPs. A) Mean CBF in the gray matter across 237 participants. B) CBF in the GM relationship with age and APOE e4 carriership. *Abbreviations:CBF = Cerebral blood flow; GM = Gray matter; APOE = Apolipoprotein E*.

## 4. Discussion and Future Directions

Here, we provided a detailed description of the EPAD MRI dataset and the semi-automatic pipeline developed to process raw multimodal multicenter imaging data from the v1500 baseline data release, and illustrate its IDP feature extraction. We proposed to use a combination of distribution-based statistics and subpopulation visual assessments to help identify low-quality images in EPAD. Finally, we described the computation of MRI summary measures (IDPs) from the core and advanced sequences and provided evidence of meaningful associations between brain phenotypes and other non-imaging metadata. We anticipate that this work may benefit both EPAD investigators by providing a structured, organized, and well-characterized neuroimaging dataset.

### 4.1 MRI Cohorts Research Data Management

The proposed pipeline for the EPAD study represents an attempt to deliver standardized data in a multicenter MRI study, in a similar way to what has been previously done in other cohorts. Among publicly available neuroimaging cohorts, extensive work on the documentation of MRI data management strategies has been recently published by two large MRI cohort studies (Sudlow et al. 2015; Bookheimer et al. 2019). These efforts entail fully automated pipelines for MRI processing, QC, and computation of IDPs from multimodal single-scanner imaging datasets (F. Alfaro-Almagro and Jenkinson 2016; Glasser et al. 2013). Compared to these studies, a unique feature of our pipeline is that it caters multiple scanners from multiple vendors and harmonizes differences between different scanner outputs to allow for automated processing and semi-automated QC. Multisite neuroimaging RDM represents a more complex procedure due to the variety of DICOM output structures from different manufacturers and scanners. The multisite Alzheimer Disease Neuroimaging Initiative (ADNI, (Mueller et al. 2005)) has focused on MRI standardization across sites, mainly during the preparation phase (Jack et al. 2008) where QC and preprocessing were done centrally and also provided extracted MRI metrics to investigators. The Preprocessed Connectome Project (Cameron et al. 2013) is a community effort to systematically process data from several available multicenter datasets including structural and functional MRI workflows and comparing outputs of different processing choices and toolboxes. As an example, the Autism Brain Imaging Data Exchange (ABIDE) dataset (Di Martino et al. 2014), which includes 1114 participants with structural (3D T1w) and rs-fMRI data from 16 sites, was processed using three different structural and four functional pipelines, and corresponding derivatives were made available. While both these studies and our EPAD pipeline managed to deal with such diverse datasets, we specifically aimed at creating a unified workflow by delivering a uniform and QCed shared outcome, promoting reproducibility, and avoiding redundancy and variability of results.

### 4.2 Quality Control

Our data-informed QC procedure was focused on selecting a handful of most informative image features in a semi-automatic fashion. On the other hand, most recent MRI studies focused on predicting the scan quality automatically using a fairly large number of potentially helpful image features through unsupervised or semi-supervised methods (F. Alfaro-Almagro and Jenkinson 2016; Pizarro et al. 2016). The application of machine learning classifiers has proved its efficiency in recent QC efforts on classifying 3D T1w image-quality from QC features distribution, both the UKBiobank (F. Alfaro-Almagro and Jenkinson 2016) and ABIDE (Esteban et al. 2017; Di Martino et al. 2014) datasets. Similarly, random forest classifiers trained on FreeSurfer QC output showed good accuracy in scanning site identification, supporting the use of multivariate approaches for QC metrics’ importance evaluation (Raamana et al. 2021). However, while these works aimed at the fully automatic prediction of image quality from unseen scans/sites, we focused on identifying a set of informative QC features as a pre-selection and guide for visual inspection. Similar semi-automated procedures have also been proposed in the literature. Previously ((Bastiani et al. 2019)), dMRI QC features were used to create individual and group reports and, similar to our work, interactively inspect automatically flagged problematic scans. MRIQC (Esteban, Gorgolewski, and Poldrack 2017) provides interactive individual reports created for straightforward low-quality image visualization. Likewise, well-established processing pipelines for different MRI modalities (Esteban et al. 2019; Mutsaerts et al. 2020) produce single-subject visual QC reports for the quality assurance of specific processing steps.

We showed a valuable pragmatic and intuitive solution for identifying problematic acquisitions and reducing the number of scans for visual inspection in large multicenter cohorts. Moreover, considering the recent efforts to relate quality metrics to the output of expert visual rating (Esteban et al., n.d.), another advantage of our QC procedure was the possibility of using statistical regression models to evaluate the informative character of single QC features and their agreement with a human visual inspection, as well as their relationship with scanning site and other participants’ demographic data. In line with previous work (Esteban et al. 2017), all QC features were strongly dependent on scanners and their sites, advocating for within-site normalization of quality metrics in multicenter MRI cohorts, as similar features’ values might be related to different quality of images from different scanners. Moreover, we also showed that demographic and clinical participants’ characteristics are related to image quality metrics. Former studies had already shown that age and sex can relate to impact the quality of structural derived measures of brain atrophy (Gilmore, Buser, and Hanson 2021), and fMRI derived functional connectivity (Hodgson et al. 2017). Together with our results, these findings suggest that scan quality might confound effects attributed to clinical variables and, consequently, that fully automated QC procedures might be more prone to exclude scans from selected groups of participants (e.g. older participants or clinical groups).

### 4.3 Image-derived Phenotypes

In agreement with current literature (Damoiseaux 2017), we showed an overall effect of age in modulating brain IDPs across different MRI modalities in the EPAD-LCS data set. However, different from more clinically oriented papers, we here only reported basic association without investigating statistical significance or the effect of covariates correction on the studied associations. Similar to our findings using FreeSurfer and CAT12 segmentations, studies on structural brain converge on a gradual loss of brain volume with advancing age, specifically in the hippocampus (Salat et al. 2004). Moreover, a similar frontal and parietal prevalence of vascular disease with older age, as demonstrated with the 3D FLAIR IDPs, has been shown in previous works (Chabriat and Jouvent 2020). Similar to our findings, a nonlinear age-related decline in resting-state fMRI networks has previously been demonstrated. In (Persson et al. 2014), while participants below 66 years showed an increase in DMN connectivity over time, while participants older than 74 years showed a decline. Additionally, in (Brier et al. 2012) early cognitive changes modulated default mode and frontoparietal network connectivity, possibly altering normal aging trajectories. As observed with the dMRI IDPs, alterations of white matter integrity are a typical sign of aging brains (Barrick et al. 2010). Previous publications agree with our result of reduced TBSS values in amyloid-positive individuals (Collij et al. 2020). Finally, A decline of CBF in the GM has been widely reported in recent studies of elderly individuals (Juttukonda et al. 2021). Previous studies on CBF with APOE genotype have shown higher brain perfusion related to worse cognitive impairment in older adults carrying the APOE e4 allele (Zlatar et al. 2016), and higher regional perfusion in e4 carriers in the left cingulate and lateral frontal and parietal regions (McKiernan et al. 2020). Moreover, similar group regional variability, showing high perfusion in the cingulate, precuneus, and frontal cortices and low perfusion in basal ganglia, has been previously reported (Pfefferbaum et al. 2010).

### 4.4 Limitations

Although we acknowledge the heterogeneity of existing MRI processing methodologies and implementations, we focused here on a purely descriptive overview of the procedures implemented for the EPAD study. As we followed generally accepted and standard pipelines, a potential limitation of this study could be the lack of more novel and AI-based processing routines. For example, performing denoising of signal drift correction is becoming a preferable procedure for dMRI preprocessing (Vos et al. 2017), even if few implementations exist. Another weakness of the proposed pipeline is the lack of longitudinal routines. Challenges of longitudinal MRI processing and QC entail the necessity of additionally taking into account within-subject variability, which could be added to this pipeline in future extensions (Mills and Tamnes 2014). Moreover, one main limitation of the validation approach used for the QC workflow is the partial circularity of constructing and testing the visual QC assessment based on the estimated QC features. Nonetheless, while our first aim was to automatically flag poor quality images for visual inspection, we then elaborated on the informative characters of QC features, studying their association with visual judgment. Furthermore, in contrast to more systematic procedures, in which visual inspection is performed on the whole sample (Waber et al. 2007), we only focused on automatically flagged scans and on a subset of non-flagged. The present work is a pragmatic approach of combining image feature-based QC with visual QC of a limited number of scans, which may be more feasible for large imaging cohorts. Reflective of EPAD’s multi-center and multi-vendor design, ExploreQC was tailored to this dataset and requires further validation and testing before it can be generalized to other studies. Eventually, the lack of visual QC standards for advanced MRI sequences motivates the development of more standardized approaches in future works. The IDPs found in the EPAD v1500.0 data sample cover a broad range of structural and functional brain phenotypes. However, several possibilities for summary brain measures exist (Gong, Beckmann, and Smith 2020). Future efforts should focus on widening the present number of IDPs to entail a new range of brain phenotypes, including measures of longitudinal changes where applicable.

## Conclusion

We provide a detailed description of the baseline EPAD LCS MRI dataset including the processing and QC procedures and details of the computation of derived data from core and advanced MRI sequences yielding biologically plausible IDPs. The introduced procedures and results may serve as a reference point for future developments and promote replicability and stability of results on the EPAD cohort. We made the pipeline available for external investigators aiming at the comparability of outcomes between different cohorts. We anticipate that this work will help both imaging and non-imaging researchers working on EPAD for an easier understanding and use of the shared data.

## List of abbreviations

ASL: Arterial spin labeling
LST: Lesion Segmentation Toolbox
BIDS: Brain Imaging Data Structure
MRI: Magnetic Resonance Imaging
CSF: Cerebrospinal Fluid
NIfTI: Neuroimaging Informatics Technology Initiative
DICOM: Digital Imaging Communication in Medicine
QC: Quality Control
dMRI: Diffusion MRI
RDM: Research Data Management
EPAD: European Prevention of Alzheimer’s Dementia
rs-fMRI: Resting-state functional MRI
EPI: Echo-planar imaging
SPM: Statistical Parametric Mapping
FLAIR: Fluid-attenuated inversion recovery
SWI: Susceptibility weighted imaging
FSL: FMRIB Software Library
TE: Echo Time
GIF: Geodesic Information Flows
TR: Repetition Time
GM: Gray Matter
WMH: White Matter Hyperintensities
IDP: Image-derived phenotype
WM: White Matter
LEAP: Learning Embedding for Atlas Propagation

## Acknowledgments

This work is part of the EPAD LCS (European Prevention of Alzheimer’s Dementia Longitudinal Cohort Study). The authors would like to express their gratitude to the EPAD-LCS participants, without whom this research would have not been possible.

