## Supplementary Materials for "The European Prevention of Alzheimer’s Dementia (EPAD) MRI Dataset and Processing Workflow"

### Supplementary material

#### 1. EPAD imaging Pipeline

| Processing Step | Software's Implementation | Specifics | Code Availability |
|---|---|---|---|
| **1. Data Curation** | | | |
| 1.1 DICOM Curation Module | ExploreASL | Harmonization of DICOM raw folder structure across sites and modalities | https://github.com/ExploreASL/ExploreASL/tree/EPAD |
| 1.2 DICOM Import Module | Dcm2niiX, ExploreASL | Converts DICOM to NIfTI | https://www.nitrc.org/projects/dcm2nii/ |
| **2. Data Pre-processing** | | | |
| 2.1 Core-sequences pre-processing | ExploreASL<br>SPM, CAT12, LST, BaMoS | Standard pre-processing of structural core sequences (3D T1w and 3D FLAIR) | https://github.com/ExploreASL/ExploreASL/tree/EPAD |
| 2.2 Advanced Sequences pre-processing | ExploreASL,<br>SPM12,<br>FSL | Standard pre-processing of resting-state functional, diffusion and ASL MRI sequences | |
| **3. Data Quality Control** | | | |
| 3.1 Feature estimation Module | ExploreQC | Computes QC features for different image modalities | https://github.com/luislorenzini/ExploreQC/tree/EPAD |
| 3.2 Visualization Module | ExploreQC | Interactively visualizes parameters distributions within and between sites | |
| **4. Image Derivatives** | | | |
| 4.1 Core Derivatives | LEAP, FreeSurfer, BaMoS | Computes GM global and local volumes and thickness, global WMH volume | NA |
| 4.2 Advanced Derivatives | FSL melodic/TBSS, ExploreASL | Computes resting-state network connectivity (rs-fMRI), global and local FA values (DTI), cerebral blood flow and spatial CoV (ASL) | NA |

**Table S1. Overview of processing steps and implementation in the EPAD imaging cohort. Adapted from** (Mutsaerts et al. 2020).Version of software used are: ExploreASL v1.0.2; SPM12 (r7771); FSL 6.0.2; FreeSurfer v6.0; *Abbreviations: DICOM = Digital imaging and communications in medicine; NIfTI = Neuroimaging informatics technology initiative; SPM = Statistical parametric mapping; FSL = FMRIB Software Library ; BaMos = Bayesian model selection; FLAIR = Fluid Attenuated Inversion Recovery; ASL = Arterial Spin Labeling; QC = Quality Control; LEAP =Learning Embeddings for Atlas Propagation; GM = Gray Matter; WM = White Matter; WMH = White matter hyperintensities; TBSS = Tract-based spatial statistics; rs-fMRI = resting-state functional MRI; DTI = Diffusion Tensor Imaging; CoV = Coefficient of variation.*

### 2. Data sharing procedure

Data is accessible upon submission of a data request through the EPAD LCS Research Access Portal (ERAP; http://ep-ad.org/erap/). To ensure consistency and comparability of results, researchers are encouraged to use pre-processed MRI data. However, both raw and derived imaging data, in the form of Neuroimaging Informatics Technology Initiative(NIfTI) files, are made available through the XNAT (https://xnat.org) imaging informatics platform (Marcus et al. 2007). DICOM files can only be shared upon an argued request. IDPs are centrally stored and shared with investigators in combination with other EPAD non-imaging metadata (see “Core/Advanced sequences derived phenotypes” sections in the manuscript). More information can be found online (http://ep-ad.org/).

### 3. DICOM Curation

<table>
<tr><th>DICOM SeriesDe</th><th>BIDS gr</th><th>BIDS ScanT</th><th>ExploreAS</th><th>Site010</th><th>Site110</th><th>Site011</th><th>Site012</th><th>Site013</th><th>Site014</th><th>Site015</th><th>Site016</th><th>Site017</th></tr>
<tr><td>pca(?!sl)</td><td>Other</td><td>PhaseContrast</td><td></td><td></td><td></td><td></td><td></td><td></td><td></td><td></td><td></td><td></td></tr>
<tr><td>(survey|scout)</td><td>Other</td><td>Survey</td><td></td><td></td><td></td><td></td><td></td><td></td><td></td><td></td><td></td><td></td></tr>
<tr><td>(derived|second</td><td>Other</td><td>Reconstructions</td><td></td><td></td><td></td><td></td><td></td><td></td><td></td><td></td><td></td><td></td></tr>
<tr><td>moco</td><td>Other</td><td>MoCo</td><td></td><td></td><td></td><td></td><td></td><td></td><td></td><td></td><td></td><td></td></tr>
<tr><td>(localizer|loc|lo</td><td>Other</td><td>Localizer</td><td></td><td></td><td></td><td></td><td></td><td></td><td></td><td></td><td></td><td></td></tr>
<tr><td>(examcard|Phoe</td><td>Other</td><td>ExamCard</td><td></td><td></td><td></td><td></td><td></td><td></td><td></td><td></td><td></td><td></td></tr>
<tr><td>(defaultpresenta</td><td>Other</td><td>ReferenceScan</td><td></td><td></td><td></td><td></td><td></td><td></td><td></td><td></td><td></td><td></td></tr>
<tr><td>(b1|b1.*cal|cal)</td><td>Other</td><td>B1Calibration</td><td></td><td></td><td></td><td></td><td></td><td></td><td></td><td></td><td></td><td></td></tr>
<tr><td>T2.*(*|star|ffe|</td><td>anat</td><td>T2star</td><td>anat_T2sta</td><td>46</td><td>47</td><td>47</td><td>47</td><td>47</td><td>47</td><td>47</td><td>47</td><td>47</td></tr>
<tr><td>(swi|swan)</td><td>swi</td><td>part_mag</td><td>swi_part_r</td><td>0</td><td>60</td><td>[60 120]</td><td>[60 120]</td><td>0</td><td>60</td><td>0</td><td>0</td><td>0</td></tr>
<tr><td>(flair|t2.*spc.*fl</td><td>anat</td><td>FLAIR</td><td>FLAIR</td><td>[192 176 2</td><td>192</td><td>192</td><td>192</td><td>192</td><td>192</td><td>192</td><td>192</td><td>192</td></tr>
<tr><td>(t2|t2w).*(?!swi</td><td>anat</td><td>T2w</td><td>anat_T2w</td><td>36</td><td>47</td><td>47</td><td>47</td><td>47</td><td>47</td><td>47</td><td>47</td><td>47</td></tr>
<tr><td>(SE-|)fMRI.*revP</td><td>func</td><td>RevPE</td><td>func_RevP</td><td>0</td><td>[38 76]</td><td>[38 76]</td><td>[38 76]</td><td>0</td><td>38</td><td>0</td><td>0</td><td>0</td></tr>
<tr><td>(SEfMRI|SE-fMRI</td><td>func</td><td>NormPE</td><td>func_Norn</td><td>0</td><td>[38 76]</td><td>[38 76]</td><td>[38 76]</td><td>0</td><td>38</td><td>0</td><td>0</td><td>0</td></tr>
<tr><td>(fmri|rsfmri|RS-</td><td>func</td><td>bold</td><td>func_bold</td><td>0</td><td>204</td><td>7752</td><td>7752</td><td>0</td><td>204</td><td>0</td><td>0</td><td>0</td></tr>
<tr><td>(B(0|o)|DTI)(-|_</td><td>dwi</td><td>RevPE</td><td>dwi_RevPE</td><td>0</td><td>[60 120 1</td><td>[60 120 1</td><td>[60 120 180</td><td>0</td><td>[55 60]</td><td>0</td><td>0</td><td>0</td></tr>
<tr><td>dti</td><td>dwi</td><td>dwi</td><td>dwi</td><td>0</td><td>[3233 324</td><td>3300</td><td></td><td>0</td><td>[55 60]</td><td>0</td><td>0</td><td>0</td></tr>
<tr><td>m(0|o).*revpe</td><td>asl</td><td>RevPE</td><td>ASL4D_Re</td><td>0</td><td>0</td><td>0</td><td>0</td><td>0</td><td>[55 60]</td><td>0</td><td>0</td><td>0</td></tr>
<tr><td>m(0|o).*TR2</td><td>asl</td><td>FitM0AndT1</td><td></td><td>0</td><td>0</td><td>0</td><td>0</td><td>0</td><td>0</td><td>0</td><td>0</td><td>0</td></tr>
<tr><td>m(0|o).*(|TR10)</td><td>asl</td><td>M0</td><td>M0</td><td>0</td><td>0</td><td>0</td><td>0</td><td>0</td><td>0</td><td>0</td><td>0</td><td>0</td></tr>
<tr><td>(pcasl|pasl|asl|</td><td>asl</td><td>asl</td><td>ASL4D</td><td>0</td><td>20</td><td>20</td><td>63</td><td>0</td><td>0</td><td>0</td><td>0</td><td>0</td></tr>
<tr><td>(t1|mprage|3d_i</td><td>anat</td><td>T1w</td><td>T1</td><td>172</td><td>176</td><td>176</td><td>176</td><td>196</td><td>176</td><td>176</td><td>176</td><td>170</td></tr>
</table>

**Figure S1. First columns of the TSV file specifying regular expression and sequence characteristics per scan.** This information is used by the DICOM curation module to unpack any zipped DICOMs, correct the folder-depth and sort DICOMs based on their scan types. This information is also used to make the data BIDS-compatible by restructuring the folder structure and assigning ‘runs’ in case of repeated scans. TSV = tab-separated values , DICOM = Digital Imaging and COmmunications in Medicine, BIDS = Brain imaging data structure.

### 4. ExploreQC: Software Specifications

ExploreQC is an extension to the MRI image processing toolbox ExploreASL (Mutsaerts et al. 2020) and as such is partially dependent on ExploreASL functions. The features estimation module is built in MATLAB 2015a. The visualization module with its interactive dashboard is written in R and based on the shiny package (https://shiny.rstudio.com/). ExploreQC functionalities are included in our pipeline and can be found in https://github.com/ExploreASL/ExploreASL/tree/EPAD. Currently, ExploreQC has been specifically developed and tested for the EPAD cohort, future work will be needed to generalize its functionalities on other datasets.

### 5. ExploreQC: Feature estimation module

Features are computed over 5 Image Features Domains describing general quality issues that can be found in MRI:

- *Motion.* Resulting from involuntary movements (e.g. respiration, cardiac motion and blood flow, eye movements and swallowing) or a change of position in the scanner (for a review (Zaitsev, Maclaren, and Herbst 2015)).
- *Noise.* Random signal variations of no interest. Thermal noise from the equipment and physiological noise from the subject generally contribute to image noise (Liu 2016).
- *Inhomogeneity.* Caused by MR coil nonuniformity and local perturbations of the main magnetic field, resulting in smooth intensity variations across the image (Peltonen, Mäkelä, and Salli 2018).
- *Asymmetry.* On the left-right axis, can be due to problems in the acquisition of the scan but also be linked to pathological conditions. In itself, asymmetry is not an acquisition artefact but it does violate assumptions for operations like standard space mapping.
- *Descriptives*. Intensity distributions in several tissue classes (e.g. gray matter) provide statistical descriptive characteristics of the data.

**Table S2. Features computed in the ExploreQC Feature estimation module.**

| Modality | Parameter | Domain | Description | Formula |
|---|---|---|---|---|
| 3D-T1w | | | | |
| | Signal To Noise Ratio (SNR) | Noise | Computed within the GM Mask. Noise is defined as standard deviation within a White Matter reference region (WMref). Higher values indicate a better image quality (Parrish et al. 2000) | $\frac{\mu(GM)}{\sigma(WMref)}$ |
| | Contrast to Noise Ratio (CNR) | Noise | Differences between SNR in the GM and the WM. Higher values indicate a better image quality ((Magnotta, Friedman, and FIRST BIRN 2006) | $\frac{\mu(GM)-\mu(WM)}{(WMref)}$ |
| | Coefficient of Joint Variation (CJV) | Inhomogeneity /Noise | Joint variation of GM and WM. Higher values relate to the presence of image inhomogeneity or heavy head motion artefacts (Ganzetti, Wenderoth, and Mantini 2016) | $\frac{\sigma(GM)-\sigma(WM)}{\mu(GM)-\mu(WM)}$ |
| | Foreground-Background Energy Ratio (FBER) | Inhomogeneity /Motion | Compares within head intensities to the ones outside the head (Shehzad et al. 2015) | $\frac{\mu(GM+WM+CSF)}{\mu(WMref)}$ |
| | Entropy Focus Criterion (EFC) | Motion | Shannon entropy of voxel intensities proportional to maximum possible entropy for similarly sized image, indicating ghosting and head motion-induced blurring (Esteban et al. 2017). | |
| | Asymmetry Index percentage (AI_perc) | Asymmetry | Index of voxel-wise asymmetry | $100\times\frac{(Left-Right)}{0.5\times(Left-Right)}$ |
| | Bias Index (BI) | Inhomogeneity | Standard deviation of the bias field, computed using SPM toolbox (Peltonen, Mäkelä, and Salli 2018) | |
| | Image Quality Rate (IQR) | Inhomogeneity /Noise | Image quality rate as computed by CAT12 (Gaser 2009): combination of noise, inhomogeneity, and resolution ratings | |
| | WM2MAX | Inhomogeneity /Descriptives | Median intensity within the WM mask over the 95% percentile of the full intensity distribution, that captures the existence of long tails due to hyper-intensity of the carotid vessels and fat (Esteban et al. 2017) | $\frac{\eta(GM+WM+CSF)}{95perc(WMref)}$ |
| | Descriptives | Descriptives | Descriptive statistics (e.g. max, mean, median, kurtosis, skewness) of intensities in different tissue-types | |
| rs-fMRI | | | | |

| | | | | |
|---|---|---|---|---|
| | Temporal Signal to Noise Ratio (tSNR) | Noise | SNR computed over time courses in the three brain tissues. Uses the standard deviation within a WM reference region (WMref) to estimate noise (Esteban et al. 2017) | $\frac{\mu(GM/WM/CSF)}{(WMref)}$ |
| | Global Correlation | Motion | Mean voxel-wise global correlation of voxel time series. Higher correlations relate to motion (Esteban et al. 2017) | |
| | Foreground-Background Energy Ratio (FBER) | Inhomogeneity /Motion | See above | $\frac{(GM+WM+CSF)}{\mu(WMref)}$ |
| | Entropy Focus Criterion (EFC) | Motion | See above | |
| | Framewise Displacement (FD) | Motion | Mean, standard deviation and maximum peak of motion as computed by SPM realign toolbox | |
| | Descriptives | Descriptives | Descriptive statistics of BOLD signal in GM | |
| dMRI | | | | |
| | Noise | Noise | Standard deviation of the noise map computed as in (Veraart et al. 2016), i.e. using random matrix theory | |
| | Sum of Squared Errors (SSE) | Inhomogeneity /Noise | Computes the mean voxel-wise SSE in the GM segmentation. Indicates the good fitting of the model to the data. Lower values are better | |
| | Motion Translation along the 3 axis | Motion | Average of absolute values of voxel-wise translation in millimetres in 3 dimensions (3 features) | |
| | Motion Rotation along the three axis | Motion | Average of absolute values of voxel-wise rotation in millimeters along the three brain axis (3 features). | |
| | Absolute Motion | Motion | Average motion with respect to a reference scan in millimetres. | |
| | Relative Motion | Motion | Average motion with respect to the previous scan. | |
| | Percentage of motion outliers | Motion | Percentage of slices classified as motion outliers and showing partial or total signal drop out. | |
| | Percentage of FA outliers | Descriptives | Percentage of voxels showing fractional anisotropy values > 1 | |
| | Descriptives | Descriptives | Descriptive statistics (e.g. max, mean, median), of fractional anisotropy and apparent diffusion coefficient in GM. | |

*Abbreviations: T1w = T1 weighted image; WMref = White matter reference region; GM = Gray matter; WM = White matter; CSF = Cerebrospinal fluid; SPM = Statistical parametric mapping; CAT12 = Computational anatomy toolbox; WM2MAX = White matter to maximum intensity ratio; rs-fMRI = resting-state functional magnetic resonance imaging; BOLD = Blood oxygenation level dependent; FA = Fractional anisotropy.*

**Table S3. Association of QC features computed on 3D T1w, fMRI and DTI images with participants demographic and clinical characteristics**

| | | Site | Age | | Sex (male) | | MMSE | | Amyloid (A+) | | APOE (E4) | |
|---|---|---|---|---|---|---|---|---|---|---|---|---|
| Modality | Parameter | p-value | Beta | p-value | Beta | p-value | Beta | p-value | Beta | p-value | Beta | p-value |
| **3D T1w** | | | | | | | | | | | | |
| | SNR | <0.001 | -0.30 | <0.001 | -0.43 | 1 | 0.25 | 0.15 | -1.19 | 0.01 | 0.8 | 0.2 |
| | CNR | <0.001 | -0.01 | <0.001 | -0.04 | 0.006 | 0.01 | 0.007 | -0.01 | 1 | -0.02 | 1 |
| | CJV | 0.36 | 0.01 | 0.12 | 0.06 | 0.21 | -0.01 | 1 | 0.02 | 1 | 0.01 | 1 |
| | FBER | <0.001 | -1.16 | <0.001 | -1.65 | 0.98 | 0.71 | 0.5 | -4.3 | 0.01 | 3.26 | 0.09 |
| | EFC | <0.001 | -0.02 | <0.001 | 0.47 | <0.001 | 0.04 | 0.003 | -0.08 | 0.2 | -0.01 | 1 |
| | Asymmetry | <0.001 | -0.06 | 1 | -3.46 | <0.001 | -0.35 | 1 | -0.72 | 1 | -.46 | 0.9 |
| | IQR | <0.001 | 0.01 | 0.01 | 0.03 | 0.06 | -0.01 | 0.06 | -0.02 | 1 | 0.02 | 1 |
| | BI | <0.001 | -0.01 | 0.04 | 0.06 | <0.001 | 0.01 | 1 | 0.01 | 1 | -0.01 | 1 |
| | GM Kurtosis | <0.001 | 0.01 | 1 | -0.01 | 1 | 0.01 | 1 | 0.01 | 1 | -0.01 | 1 |
| | WM Kurtosis | <0.001 | 0.01 | 0.002 | 0.03 | 1 | -0.05 | 0.01 | 0.09 | 0.7 | -0.01 | 1 |
| | CSF Kurtosis | <0.001 | 0.02 | 1 | 0.02 | 1 | 0.01 | 1 | -0.14 | 0.02 | 0.01 | 1 |
| | WM2MAX | <0.001 | -0.01 | 1 | -0.01 | 0.08 | -0.01 | 1 | -0.01 | 1 | -0.01 | 1 |
| **fMRI** | | | | | | | | | | | | |
| | GM tSNR | <0.001 | -0.14 | 0.14 | -0.11 | 1 | 0.32 | 1 | 0.46 | 1 | -0.43 | 1 |

| | | | | | | | | | | | | |
|---|---|---|---|---|---|---|---|---|---|---|---|---|
| | WM tSNR | <0.001 | -0.13 | 0.29 | -0.70 | 1 | 0.29 | 1 | 0.25 | 1 | -0.11 | 1 |
| | CSF tSNR | 0.003 | 0.05 | 1 | 1.3 | 0.03 | 0.21 | 1 | 0.73 | 1 | -0.46 | 1 |
| | tSNR GM-WM | <0.001 | 0.005 | 0.08 | -0.30 | 1 | 0.20 | 1 | 0.16 | 1 | -0.18 | 1 |
| | Motion mean | <0.001 | 0.001 | 0.001 | 0.008 | 1 | -0.002 | 1 | -0.004 | 1 | 0.003 | 1 |
| | Motion SD | <0.001 | 0.001 | 0.39 | 0.009 | 0.9 | -0.002 | 1 | -0.001 | 1 | 0.003 | 1 |
| | Motion Max | <0.001 | 0.004 | 1 | 0.07 | 0.86 | -0.01 | 1 | -0.008 | 1 | 0.04 | 1 |
| | GC | <0.001 | -0.001 | 1 | 0.001 | 1 | 0.001 | 1 | 0.004 | 1 | -0.001 | 1 |
| | FBER | <0.001 | -0.78 | 1 | 65.88 | 1 | 18.7 | 1 | 54.4 | 1 | 17.14 | 1 |
| | EFC | <0.001 | -1.96 | <0.001 | 41.18 | <0.001 | 0.04 | 0.34 | -0.08 | 1 | 0.008 | 1 |
| | Ghost to Signal | <0.001 | -0.001 | 1 | -0.006 | <0.001 | -0.001 | 1 | 0.002 | 0.09 | -0.001 | 1 |
| | GM mean | <0.001 | -623.9 | 1 | -18448.06 | 0.54 | 2384 | 1 | 22459.8 | 0.36 | -103.8 | 1 |
| | GM median | <0.001 | -651.8 | 1 | -18898.36 | 0.52 | 2392 | 1 | 22752 | 0.36 | 209.5 | 1 |
| **dMRI** | | | | | | | | | | | | |
| | Translation x | 1 | 0.002 | 1 | -0.02 | 1 | -0.002 | 1 | -0.01 | 1 | -0.01 | 1 |
| | Translation y | <0.001 | -0.005 | 0.17 | 0.05 | 1 | 0.01 | 1 | 0.01 | 1 | 0.03 | 1 |
| | Translation z | 1 | 0.001 | 1 | 0.006 | 1 | -0.005 | 1 | 0.04 | 1 | -0.04 | 0.77 |
| | Rotation x | 1 | -0.001 | 1 | -0.001 | 1 | -0.001 | 1 | 0.001 | 1 | -0.001 | 1 |
| | Rotation y | 1 | -0.001 | 1 | -0.001 | 1 | -0.001 | 1 | 0.001 | 1 | 0.001 | 1 |
| | Rotation z | 1 | 0.001 | 1 | -0.001 | 1 | 0.001 | 0.85 | 0.001 | 1 | 0.001 | 1 |
| | Absolute motion | <0.001 | 0.001 | 1 | -0.02 | 1 | 0.007 | 1 | -0.01 | 1 | 0.001 | 1 |
| | Relative Motion | <0.001 | 0.001 | 0.4 | -0.002 | 1 | -0.003 | 1 | -0.008 | 1 | -0.003 | 1 |
| | Motion outliers | <0.001 | 0.002 | 1 | 0.04 | 1 | 0.05 | 1 | -0.02 | 1 | 0.05 | 0.75 |
| | Induced distortion | <0.001 | -0.001 | 1 | -0.001 | 1 | -0.001 | 1 | 0.002 | 1 | -0.001 | 1 |
| | Noise SD | <0.001 | -0.1 | 1 | -50.74 | 1 | -5.47 | 1 | 69.46 | 0.47 | -5.47 | 1 |
| | SSE | <0.001 | -0.001 | 1 | 0.11 | <0.001 | 0.009 | 1 | -0.02 | 1 | 0.009 | 1 |

For each QC feature for fMRI and DTI, the output of a linear model (beta coefficients and p values) investigating the effect of site, age and sex is reported. Site was considered as a dummy variable and has therefore no estimated beta coefficient. *Abbreviations: GM = Gray Matter; WM = White Matter; CSF = Cerebrospinal Fluid.*

### 6. Image-Derived Phenotypes

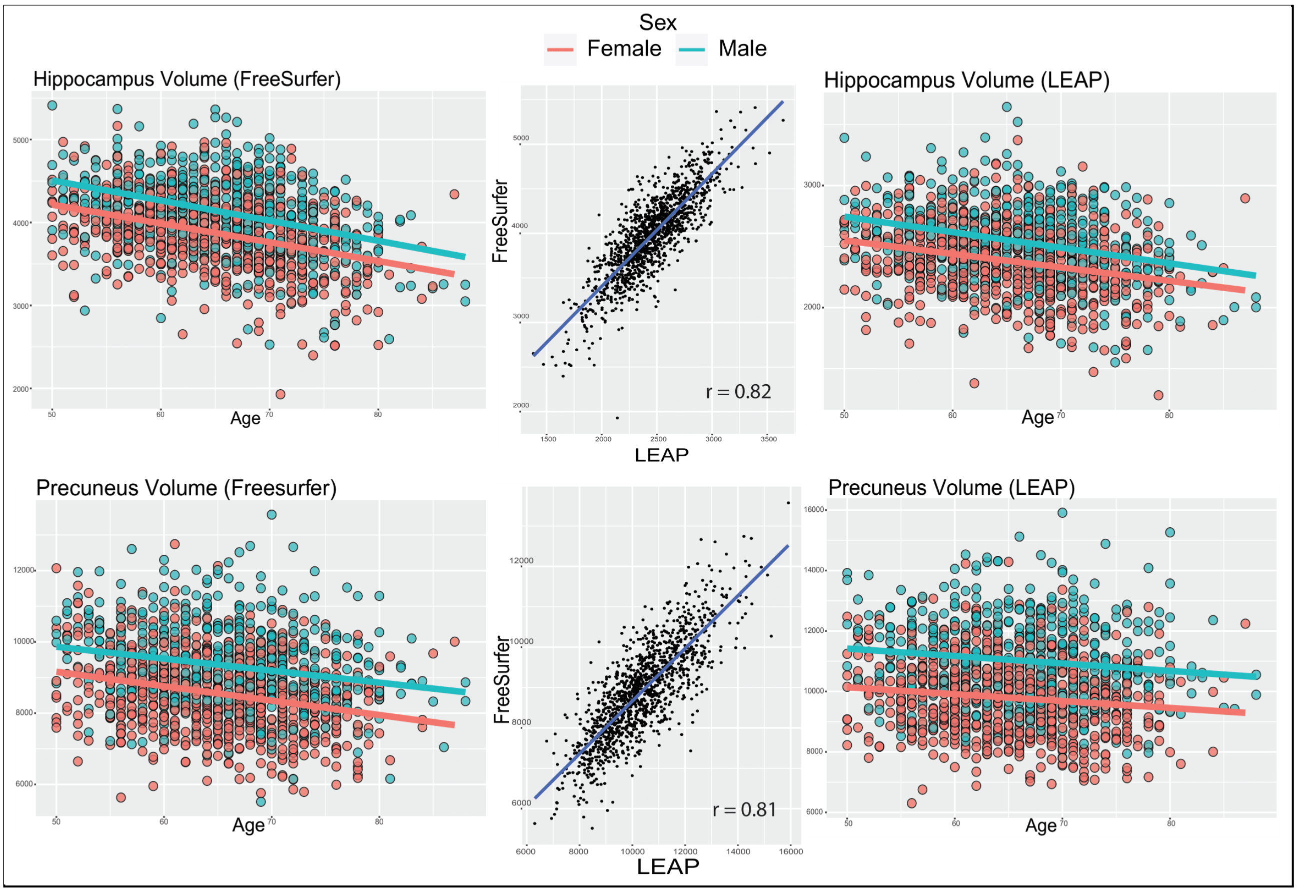


**Figure S2. LEAP and FreeSurfer derived data.** *Left:* FreeSurfer volumes for Hippocampus (top) and Precuneus (bottom) association with age and sex; *Right:* LEAP volumes for Hippocampus (top) and Precuneus (bottom) association with age and sex; *Middle:* Relationship between segmented volumes from the two pipelines. *Abbreviations: r = Pearson correlation coefficient, LEAP = Learning Embeddings for Atlas Propagation.*

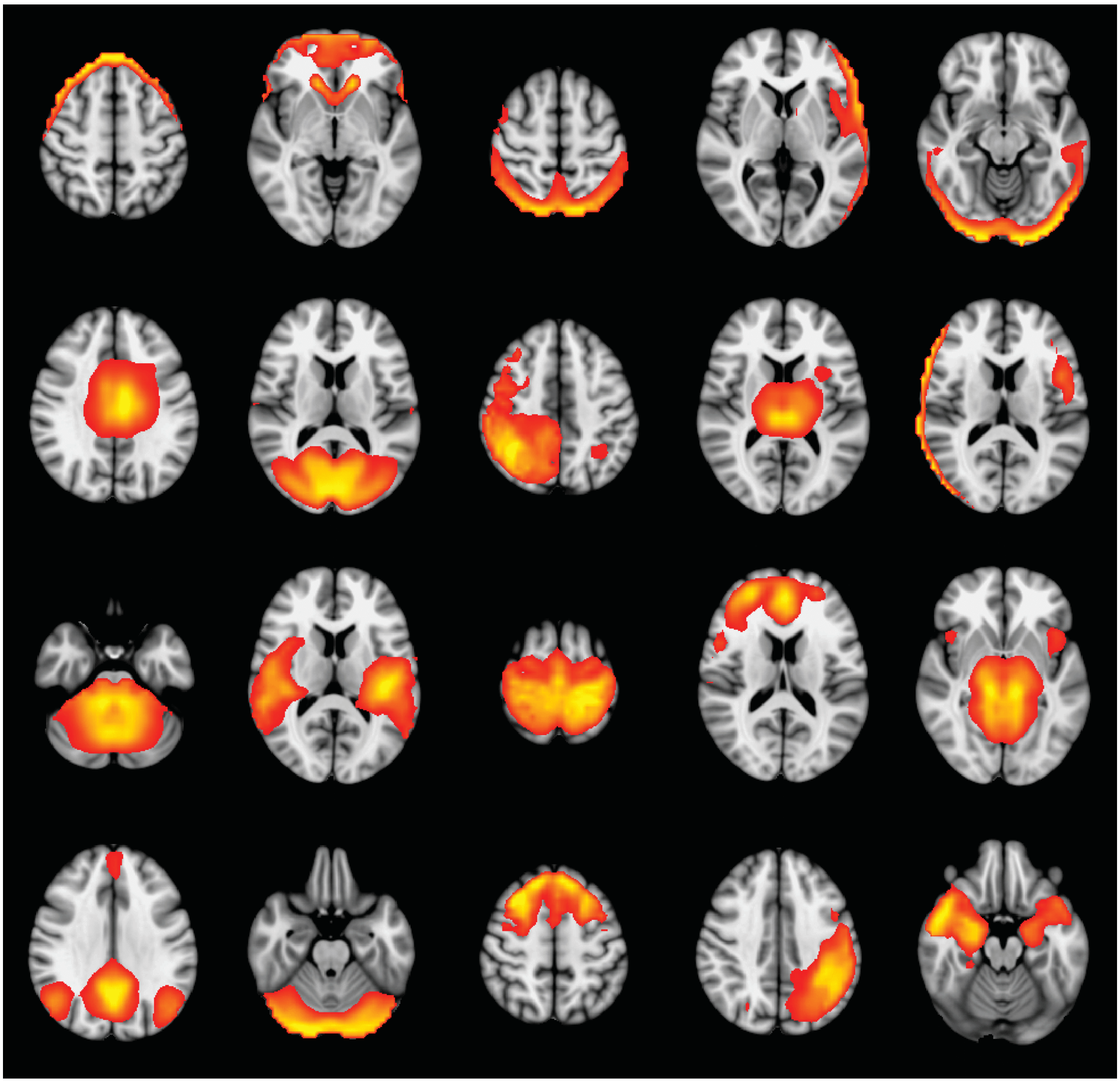

**Figure S3. Low Dimensional ICA on rs-fMRI.** 20 Resting-state networks spatial maps computed with FSL melodic. *Abbreviations: ICA = Independent component analysis.*

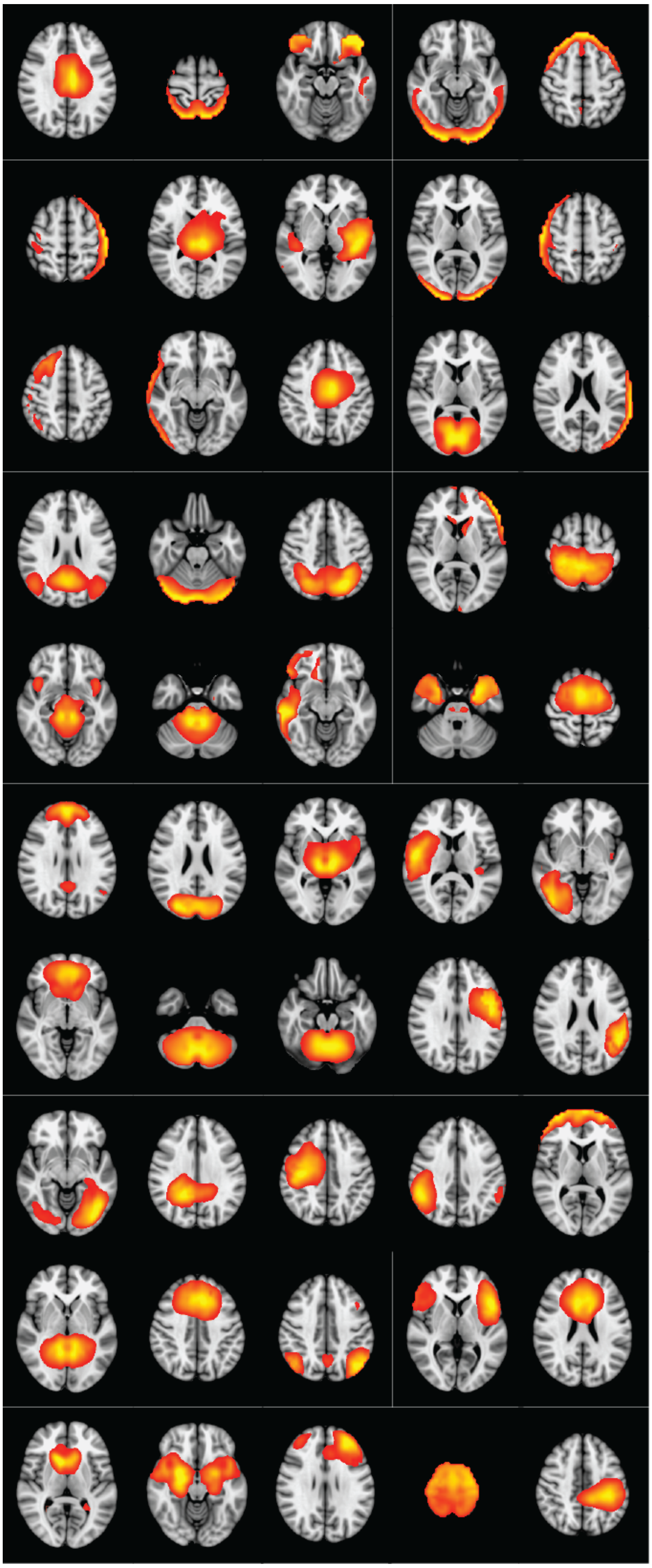

**Figure S4. High Dimensional ICA on rs-fMRI.** 50 Resting-state networks maps computed with FSL melodic. *Abbreviations: ICA = Independent component analysis*.
